# Developmental rewiring of EHMT1 function during human cortical neurogenesis

**DOI:** 10.64898/2026.10.09.757098

**Authors:** Kritika Kasliwal, Kaushiki Chatterjee, Vittoria Dickinson Bocchi, Syed R. Mahmood, Christopher M Uyehara, William Marchese, Alejandra Laguillo-Diego, Emily Goetzler, Gabriel Theis, Andrew Minotti, Ryan Walsh, Alexander Polyzos, Eftychia Apostolou, Simone Sidoli, Lorenz Studer, Matthias Stadtfeld

## Abstract

Ubiquitously expressed chromatin regulators orchestrate cell-type-specific gene expression programs, yet how they acquire distinct regulatory functions during development remains fundamentally unresolved^1,2^. Using temporally controlled protein depletion during human cortical neurogenesis *in vitro*, we show that the autism-associated histone methyltransferase EHMT1^3,4^ is developmentally repurposed from epiblast to cortical neurons. Broadly expressed zinc finger proteins recruit EHMT1 to distinct cis-regulatory elements within changing chromatin landscapes associated with lineage-specific transcription factor activity, with extensive remodeling of EHMT1 enhancer occupancy during neural induction dependent on ZNF462. This genomic rewiring is accompanied by a fundamental shift from restraining premature developmental gene expression in pluripotent cells to supporting neuronal programs, with distinct transcriptional consequences of EHMT1 loss across neural progenitors and excitatory and inhibitory neuronal subtypes. Timed depletion and recovery experiments identify the neural progenitor stage as the onset of a developmental period during which EHMT1 is required for faithful neuronal maturation. Remarkably, EHMT1 can re-engage stage-specific targets after prolonged deficiency, allowing extensive transcriptional recovery with only limited molecular scarring. These findings reveal how ubiquitous epigenetic machinery acquires developmental specificity while demonstrating the resilience of developmental gene regulatory programs to prolonged disruption, with implications for neurodevelopmental disorders.

## Introduction

Controlled remodeling of chromatin activity at individual gene loci and larger genomic domains is essential for establishing cell-type-specific gene expression programs during organogenesis^1^. Despite their often broad expression across tissues and developmental stages, chromatin regulators orchestrate highly specialized regulatory programs in distinct cell types^2^, yet the mechanisms by which they acquire these stage– and cell-type-specific functions remain poorly understood. Human cortical development provides a striking example of this challenge, as neuronal specification and maturation require exquisitely coordinated chromatin regulation over an extended developmental timeline^5,6^. Consistent with this central role, mutations in chromatin regulators frequently cause neurodevelopmental disorders^7–9^, with mutations in distinct chromatin regulators underlying neurodevelopmental chromatinopathies converging on shared transcriptional programs, cellular phenotypes, and developmental mechanisms, despite retaining disorder-specific molecular features^10,11^. This underscores the importance of understanding how ubiquitously expressed chromatin regulators acquire cell-type-specific regulatory functions during development.

Despite their developmental and clinical relevance, fundamental questions about the molecular regulation of chromatin modifiers remain unanswered. These include how distinct recruitment mechanisms are integrated to direct specific chromatin modifiers to their genomic targets in a stage– and cell-type-specific manner^12–14^, and how such stage-specific targeting determines why only a restricted subset of genes is transcriptionally sensitive to loss of individual modifiers^15^. In addition, the limited accessibility of the developing human brain has prevented a thorough characterization of epigenetic mechanisms underlying neurodevelopmental disorders and the developmental stages or cell types in which they act.

An important example of these unresolved questions is euchromatic histone lysine methyltransferase 1 (EHMT1), which was initially identified as a transcriptional repressor in naïve murine pluripotent stem cells^3^. Together with its heterodimeric partner EHMT2, EHMT1 is the primary source of the histone 3 lysine 9 dimethylation (H3K9me2) mark in euchromatic genomic regions. Most H3K9me2 is organized into broad chromatin domains that span large portions of the genome and are thought to stabilize lineage-specific gene expression programs^16,17^. Consistent with an important developmental role, studies in animal models have implicated EHMT1 in cognitive function and adaptive behavior^18,19^, while clinical studies have established EHMT1 haploinsufficiency as the cause of the autism spectrum disorder (ASD) Kleefstra syndrome^4^. Although altered gene expression has been reported in neural progenitor cells (NPCs) and cortical neurons derived from patients with Kleefstra syndrome^20,21^, how EHMT1 target selection and regulatory functions change during human corticogenesis remain largely unknown. Moreover, transient EHMT1 inhibition induces persistent developmental abnormalities during murine pluripotency acquisition^22^, raising the broader question of whether developmental EHMT1 deficiency irreversibly alters stage-specific human gene regulatory programs or whether these programs retain the capacity for subsequent molecular recovery.

Here, we use temporally controlled protein depletion during human in vitro cortical neurogenesis to determine how the autism-associated chromatin regulator EHMT1 acquires distinct regulatory functions during brain development. We show that EHMT1 is developmentally repurposed to distinct cis-regulatory elements across successive stages of cortical neurogenesis through interactions with broadly expressed zinc finger proteins whose genomic targeting is remodeled within changing lineage-specific transcription factor landscapes. This developmental rewiring shapes stage-specific transcriptional programs in neural progenitors and across multiple excitatory and inhibitory neuronal subtypes, while identifying the neural progenitor stage as the onset of a developmental period during which EHMT1 activity is required for faithful neuronal maturation. Finally, we demonstrate that despite prolonged developmental deficiency, EHMT1 retains the capacity to re-engage stage-specific targets and restore most transcriptional programs, revealing an unexpected degree of molecular resilience.

## Results

### EHMT1 has distinct stage-specific transcriptional targets during human cortical neurogenesis

To study the transcriptional effects of EHMT1 depletion at consecutive stages of cortical development in a temporally controlled manner, we used the dTAG degradation system^23^. Specifically, we targeted both alleles of the endogenous *EHMT1* locus in MEL-1 human embryonic stem cells (ESCs) with a multi-purpose cassette containing FKBP12^F36V^ for rapid and controlled protein depletion, hemagglutinin (HA) tags for efficient immunoprecipitation, and an EGFP transcriptional reporter (**Figure 1A, S1A**). Exposure of EHMT1-dTAG ESCs to the heterobifunctional degrader dTAGv1^24^ (“dTAG”) resulted in rapid (<1 h) and efficient EHMT1 depletion, followed by reduction of global H3K9me2, which – similar to reports in other systems^25,26^ – fell to about 40% of starting levels after 48 h (**Figure S1B**).

**Figure 1.**
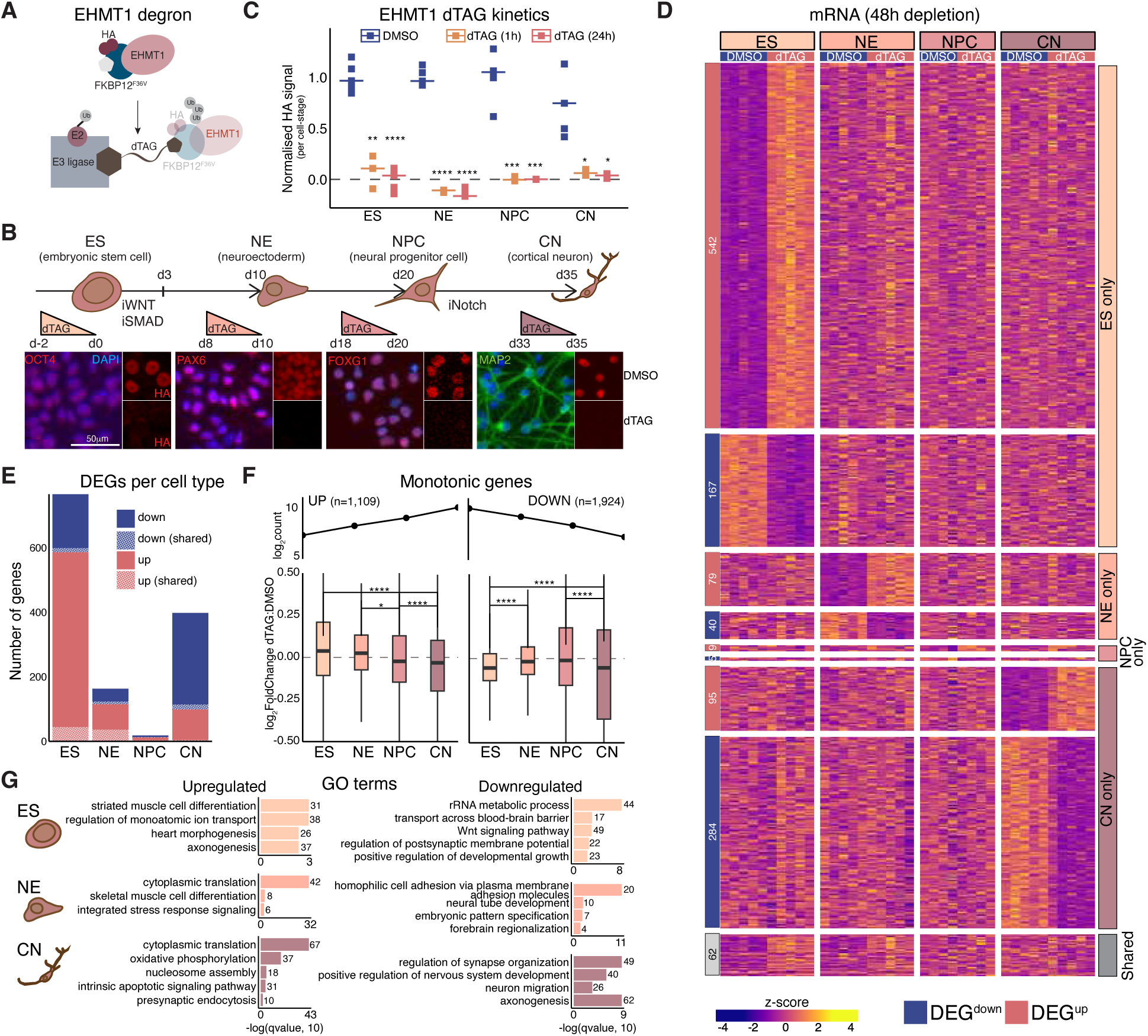
EHMT1 has distinct stage-specific transcriptional targets during human cortical neurogenesis. **A**, Schematic of dTAG-induced depletion of the EHMT1-HA-FKBP12^F36V^ fusion protein. **B**, Paradigm of human cortical neuronal differentiation, with immunofluorescence (IF) validation of stage-specific markers and dTAG-induced depletion of EHMT1(HA). **C**, IF quantification of EHMT1 depletion kinetics at 1 and 24 h post-dTAG treatment across ES, NE, NPC, and CN stages for one representative clone (n=5 cells per condition). Signal was compared to DMSO using an unpaired t-test (****p < 0.0001, ***p < 0.001, **p < 0.01, *p < 0.05, n.s. – ‘not significant’. Note: the same significance labels apply throughout. **D**, Heatmap of differentially expressed genes (DEGs; padj < 0.05, |log2FoldChange| > 0.5) following 48 h of EHMT1 depletion at four stages of neurogenesis. Z-scores were calculated for each cell type, and cells were clustered using stage-specific DEGs (biological clones n=5 for ES, NE, and CN; n=4 for NPC). **E,** Bar plot of total unique and shared DEGs at each stage of neurogenesis. **F,** Log2(DESeq2 normalized counts) and log2FoldChange at each neurogenesis stage for genes with strictly monotonic expression changes – increasing (left) or decreasing (right) from ES to CN. Counts were compared between consecutive stages using a paired t-test. **G,** EnrichGO Biological Processes ontology terms of genes up-or down-regulated at each stage of neurogenesis (padj < 0.05, |log2FoldChange| > 0).

We differentiated EHMT1-dTAG ESCs into cortical neurons (CNs) using a protocol that leverages timed inhibition of WNT, SMAD, and NOTCH signaling to drive cortical neurogenesis via defined intermediate stages of anterior neuroectoderm (NE) and neural progenitor cells (NPC)^27^ (**Figure 1B**). We confirmed synchronized differentiation based on the expression of stage-specific marker genes at the protein and RNA levels (**Figure 1B, S1C,D**). Importantly, EHMT1 was ubiquitously expressed throughout neurogenesis and efficiently degraded after 1 h of dTAG treatment at all stages (**Figure 1B,C, S1E**). Using this system, we transcriptionally profiled cells at four key stages of neurogenesis (ESC, NE, NPC, and CN) after 48 h of EHMT1 depletion (**Figure 1B**). Principal component analysis confirmed progressive identity changes and a high degree of reproducibility between cells derived from different (n = 4 or 5) clonal ESC lines **(Figure S1F**). Differential gene expression analysis (padj<0.05; |log2FC|>0.5) revealed dysregulation across all stages, albeit to strikingly different extents (**Figure 1D,E, S1G, Supplementary Table 1**). ESCs showed the strongest response, with genes being largely upregulated upon EHMT1 loss (586 DEG^up^ compared to 180 DEG^down^; **Figure S1G**) in agreement with the canonical repressor function of EHMT1^3^. Of note, these genes were not affected by dTAG treatment of ESCs engineered to express an EGFP-dTAG fusion (**Figure S1A, S1H**), ruling out a nonspecific transcriptional effect of engaging E3 ligases in this manner. Like ESCs, NE and NPCs predominantly responded to EHMT1 loss with transcriptional de-repression, but this effect was subdued in NE (115 DEG^up^ and 48 DEG^down^) and barely detectable in NPCs (12 DEG^up^ and 6 DEG^down^) (**Figure 1D,E, S1G**). Contrary to this trend, EHMT1 depletion in CNs resulted in several hundred DEGs, most of which were downregulated (99 DEG^up^ and 300 DEG^down^) (**Figure 1D,E, S1G**). Overall, the magnitude of the transcriptional response tracked with the extent of H3K9me2 loss across stages (**Figure S1I**), consistent with a contribution of EHMT1-dependent H3K9 methylation to these transcriptional effects.

Beyond differences in response magnitude to acute EHMT1 loss, the identity of affected genes was also highly cell-type specific. The overlap of DEGs shared between stages was limited, with only 62 of 1,284 genes (4.8%) affected in two or more cell types (**Figure 1E, S1J**). Z-score representation further highlighted the stage-specific nature of transcriptional responses (**Figure 1D**). Despite this limited overlap at the level of individual genes, several coherent developmental trends emerged. Genes whose expression increased monotonically across differentiation were, on average, precociously upregulated after EHMT1 loss in ESCs but downregulated after EHMT1 loss in cortical neurons (**Figure 1F**), indicating that EHMT1 contributes differently to the temporal execution of the cortical neurogenesis program at distinct stages. In agreement, gene ontology (GO) analysis of genes significantly affected (padj<0.05) by EHMT1 loss showed premature upregulation of differentiation-associated programs, including axonogenesis, in ESCs, downregulation of regionalization and axial patterning programs in NE, and downregulation of both neurogenesis and neuronal function-associated programs in CNs (**Figure 1G**). In addition, EHMT1-depleted NE and CN showed upregulation of stress-related programs, suggesting that EHMT1 loss also perturbs cellular programs not directly linked to differentiation. Overall, stage-specific depletion shows that the ubiquitously expressed EHMT1 regulates largely distinct transcriptional programs at successive stages of cortical neurogenesis, revealing extensive developmental rewiring of its regulatory function.

### Molecular determinants of transcriptional sensitivity to EHMT1 loss

The remarkable stage-specificity of EHMT1-responsive genes raises the question of what renders loci transcriptionally dependent on EHMT1. We addressed this question by generating data on EHMT1 genome binding (via HA ChIP-seq) and H3K9me2 distribution (via spike-in-normalized, ultra-low-input native ChIP-seq; ULI-NChIP^28^) and integrating them with the transcriptional changes observed upon EHMT1 loss. We focused on ESCs, as they exhibited the strongest transcriptional response to EHMT1 loss after acute depletion (see **Figure 1D**). EHMT1-HA ChIP-seq was highly reproducible across three clonal lines, with correlation coefficients >0.9 (**Figure S2A**). We identified >40,000 peaks (**Figure S2B**), demonstrating extensive EHMT1 engagement across the genome, with enrichment at putative gene regulatory elements, including promoters and distal enhancers (**Figure S2C,D**). ULI-NChIP (n = 2 clones) indicated a global loss of H3K9me2 levels of about 40-50%, concordant with flow cytometric quantification (**Figure S2E**). Together with our RNA-seq analysis following acute EHMT1 depletion, these datasets therefore appear suitable for identifying genomic features associated with transcriptional sensitivity to EHMT1 loss.

EHMT1 has been linked to the establishment of repressive domains^17^. Accordingly, we identified genomic blocks with elevated H3K9me2 levels in our ULI-NChIP data, confirming prior reports that ∼70% of the epigenome is organized in this manner (**Figure S2F**). Visual inspection of our data revealed broad H3K9me2 domains, while EHMT1 binding was constrained to narrow peaks (**Figure 2A, S2G**). Most gene loci upregulated upon EHMT1 depletion (DEG^up^) were located within an H3K9me2 domain and harbored EHMT1 peaks in promoter-proximal or more distal regions. This is exemplified by the locus encoding the Vitamin D receptor (*VDR*) (**Figure 2A**) and the locus encoding a subunit of the GABA type A receptor (*GABRP*) (**Figure S2G**), two proteins important for brain development and function^29,30^. Specifically, about 80% of DEG^up^ (and less than 50% of control genes) were present within H3K9me2 domains (**Figure 2B**), and these genes were significantly enriched for both proximal and distal ChIPseeker EHMT1 peaks relative to DEG^down^ and unaffected genes (**Figure 2C**). Together, these observations indicate that localization within H3K9me2 domains and local EHMT1 recruitment are defining features of transcriptionally responsive loci (**Figure 2D**). In contrast to naïve mouse pluripotent cell^31^, DEG^up^ were not organized in large genomic clusters of co-regulated genes, and no H3K9me2 domain in human ESCs harbored more than three genes derepressed upon EHMT1 depletion (**Figure 2A, S2H**).

**Figure 2.**
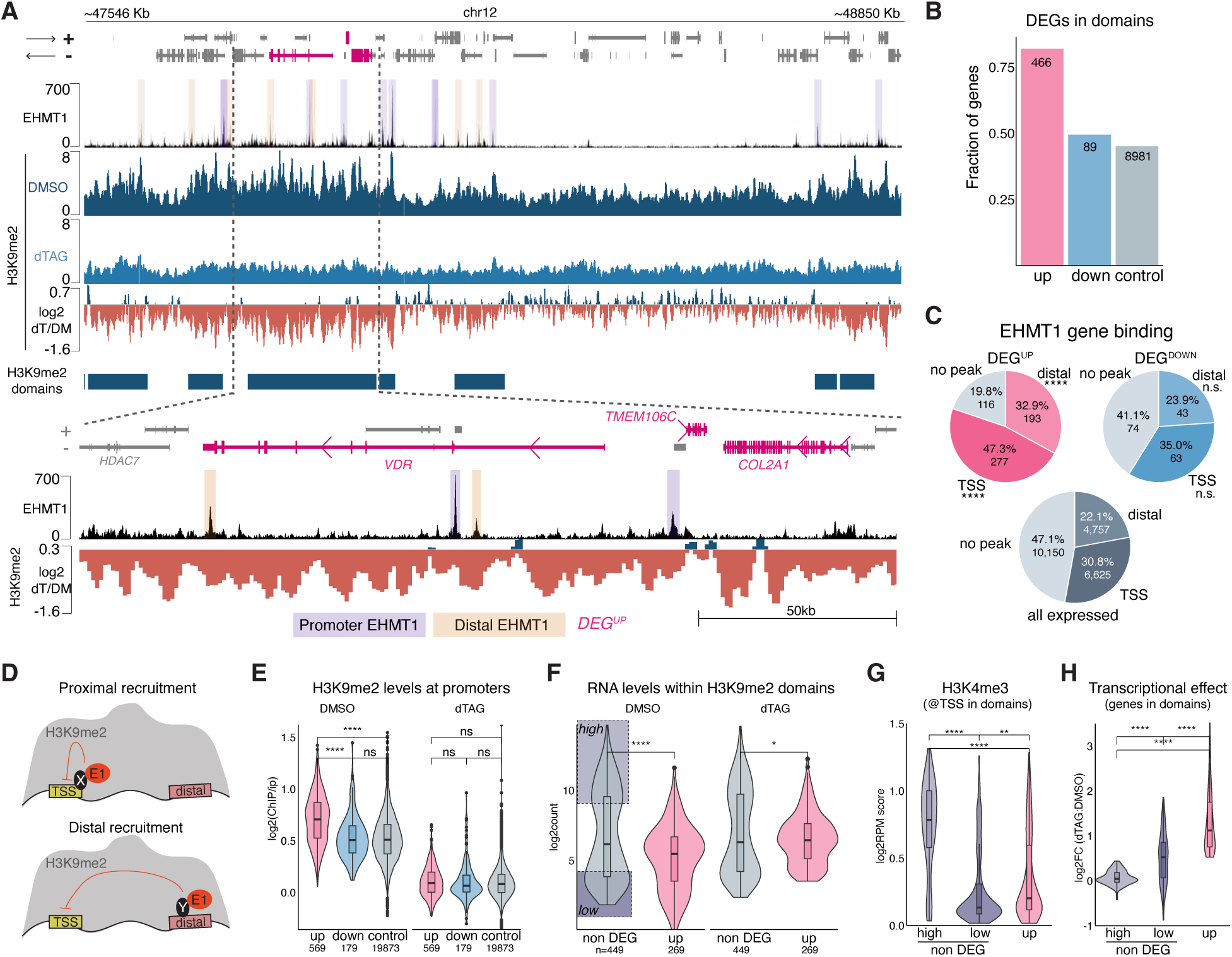
Molecular determinants of transcriptional sensitivity to EHMT1 loss. **A**, Representative ESC ChIP-seq browser tracks showing EHMT1 occupancy, EHMT1 peaks called at promoters (purple shading) or non-promoters (light brown shading), 10-kb-binned H3K9me2 signal under DMSO and 48 h dTAG treatment, the log2(dTAG/DMSO) ratio, and called H3K9me2 domains. Zoom-in panels highlight genes upregulated upon EHMT1 depletion — *VDR*, *TMEM106C*, *COL2A1*. DEGs are marked in magenta, non-DEGs in gray. **B**, Fraction of DEG categories (upregulated, downregulated, and expressed genes) located within H3K9me2 domains in ESCs. Expressed genes were defined as those with a non-NA DESeq2 adjusted p-value. **C,** ChIPseeker EHMT1 DEG category fraction annotations based on EHMT1 peak location, with promoter annotation prioritized over distal peaks, Pearson’s Chi-squared test with Yates’ continuity correction. **D,** Model of proximal and distal gene regulation by EHMT1 within H3K9me2 domains. **E,** Violin plots of H3K9me2 log2(ChIP/input) levels under DMSO and 48 h dTAG treatment, by EHMT1 DEG category, unpaired t-test. **F,** DESeq2-normalized counts of indicated gene categories subset to those containing an upregulated EHMT1 DEG within the same H3K9me2 domain as a non-DEG; genes in the top and bottom 30% of counts were labeled as “high” and “low” expressers, respectively, unpaired t-test. **G,** Signal of the active H3K4me3 mark (GSM2029350) at the transcriptional start site (TSS) of indicated gene categories, unpaired t-test. **H,** Gene expression change (DESeq2 log2FoldChange) in ES cells of indicated gene loci upon EHMT1 depletion, unpaired t-test.

Surprisingly, nearly half of all transcriptionally unaffected genes were also located within EHMT1-dependent H3K9me2 domains and associated with EHMT1 peaks (**Figure 2B,C**), suggesting additional molecular determinants render specific loci sensitive. Indeed, DEG^up^ exhibited elevated H3K9me2 levels at their transcriptional start site (TSS), which dropped to the same level upon EHMT1 depletion as at unaffected genes (**Figure 2E**). This suggests a thresholding effect, in which genomic loci with higher H3K9me2 levels depend more on EHMT1 for establishment and/or maintenance of transcriptional repression. The higher H3K9me2 levels of DEG^up^ might be linked to their localization to specific nuclear regions, as suggested by elevated LAMINB1 levels at their promoters (**Figure S2I**), consistent with observations in mouse ESCs^32^. To further elucidate molecular determinants of transcriptional derepression upon EHMT1 loss, we restricted our analysis to those H3K9me2 domains that contained at least one non-affected gene in addition to one or more DEG^up^ (n=217) (**Figure S2H**). These analyses suggested that transcriptionally insensitive genes fall into two fundamentally different classes. One category of non-responders was expressed at higher levels in steady-state than DEG^up^ (**Figure 2F**; “non-DEG^high^”) and exhibited elevated levels of the activation markers H3K4me3 and H3K27ac at their promoters (**Figure 2G, S2J**). This indicates that their local activation status shields these genes from the repressive effects of the surrounding H3K9me2 domain, rendering them transcriptionally unresponsive to EHMT1 loss (**Figure 2H**). The other category of non-responders was expressed at lower levels (**Figure 2F**; “non-DEG^low^”) and showed reduced H3K4me3 (**Figure 2G**), as well as only a muted transcriptional response to EHMT1 depletion (**Figure 2H**). However, non-DEG^low^ and DEG^UP^ exhibited similar levels of repressive chromatin marks (H3K27me3, H3K9me2, H3K9me3) (**Figure S2J**). This suggests a degree of derepression upon removal of H3K9me2 that is insufficient for stronger activation, perhaps because required activators are absent. Together, these findings support a two-layer model of EHMT1-dependent gene regulation. EHMT1 recruitment and elevated H3K9me2 define loci competent for repression, whereas local chromatin activity and the availability of activating transcription factors determine whether those loci are transcriptionally sensitive to EHMT1 loss.

### Target-specific recruitment of EHMT1 by distinct zinc finger proteins

The enrichment of EHMT1 near DEG^up^ in ESCs (**Figure 2E**) indicated that local recruitment is an important determinant of cell type-specific target gene regulation. Because EHMT1/2 lack DNA-binding domains^3^, we asked which proteins might mediate their recruitment or retention on chromatin during human cortical development. Immunoprecipitation mass spectrometry (IP-MS) across all four stages of neural development (n = 3 replicates) revealed 35 EHMT1 interaction partners (**Figure 3A, Supplementary Table 2**), with overall little correlation between whether a protein was identified as an interactor in a given cell type and its mRNA abundance (**Figure 3A**). EHMT1 interaction partners represented different functional categories, including chromatin components, RNA biology, and signaling regulators (**Figure 3B**). As expected, EHMT2 was the strongest EHMT1 interactor and was identified at all stages. Other strong interactors found in multiple cell types included the H3K9me2/3 readers CBX1 and CBX3, which can stabilize histone methyltransferases and promote heterochromatin propagation^33^, and the zinc finger transcription factors WIZ, ZNF462, and ZNF644, which in mouse cells have been shown to form a complex with EHMT1 or recruit EHMT2 to chromatin^34,35^. Thus, EHMT1 associates with a relatively stable set of broadly expressed zinc finger proteins throughout cortical neurogenesis, but not with obvious lineage-instructing transcription factors (**Figure 3A**).

**Figure 3.**
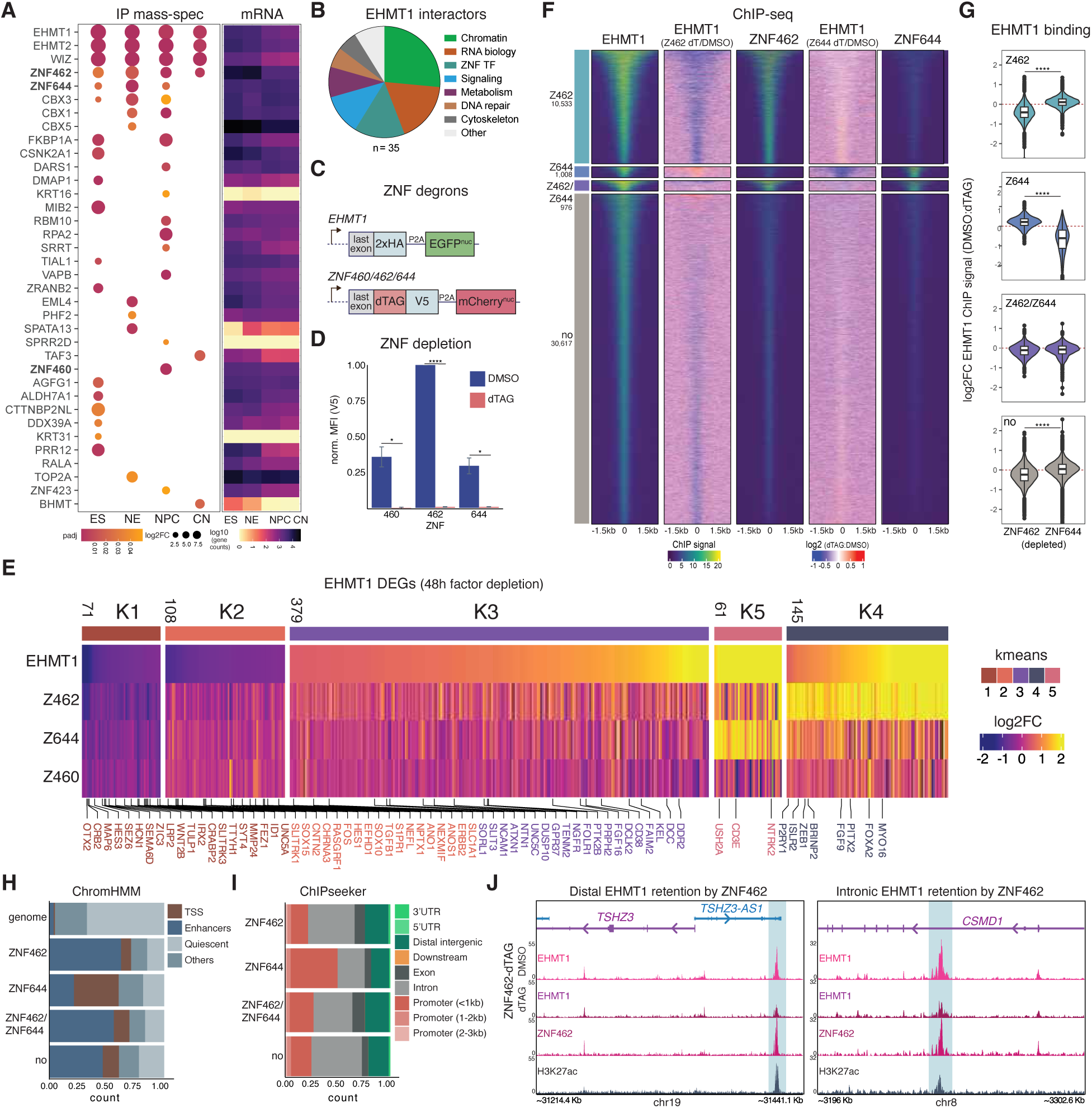
Target-specific recruitment of EHMT1 by distinct zinc finger proteins. **A**, Immunoprecipitation mass-spec (IP-MS) for EHMT1(HA) at different stages of neurogenesis (paired t-test, p < 0.05; log2FoldChange (DMSO/dTAG) > 1). Gene expression of IP-MS hits at each stage (mRNA DESeq2-normalized counts) is shown on the right. **B,** Pie chart showing functional assignments of IP-MS interactors from panel A. **C,** Simplified schematic of EHMT1-HA-GFP and ZNF-V5-dTAG-mCherry dual-tagged lines, generated for ZNF460, ZNF462, and ZNF644. **D,** Flow cytometry quantification of the indicated V5-tagged ZNFs following 48 h of dTAG-induced depletion, normalized to the maximum value (n = 3 clones, paired t-test. **E,** k-means clustered heatmap showing expression changes (dTAG/DMSO; average log2FoldChange, n = 3 clones) of EHMT1 ES cell DEGs following 48 h dTAG treatment of indicated ZNF degron lines. Genes annotated to the GO term GO:0022008 (neurogenesis) are highlighted and colored by k-means cluster. **F,** k-means clustering of EHMT1 peaks with ZNF462 and ZNF644 peaks in ESCs, as well as log2FoldChange (dTAG/DMSO) of EHMT1 peaks following depletion of ZNF462 or ZNF644. **G,** Changes in the intensity of EHMT1 signal (dTAG/DMSO; log2FoldChange) of EHMT1 peaks in each k-means cluster after ZNF462 or ZNF644 depletion, unpaired Wilcoxon test. **H,** Simplified ChromHMM 18-state chromatin model (see Fig.S2D) of the ES genome, with annotation of ChIP k-means clusters. **I,** ChIPseeker annotation of ChIP k-means clusters. **J,** Genome tracks of example NDD-associated EHMT1 ES DEG^up^ (TSHZ3, CSMD1) with EHMT1 binding mediated by ZNF462. Tracks show EHMT1 signal under ZNF462 DMSO and dTAG conditions, along with ZNF462 and H3K27ac signal.

We focused on three ZNF interactors: ZNF462, which is a category 1 SFARI gene and the cause of Weiss-Kruszka syndrome^36^, ZNF644, which has recently been associated with ASD^37^ and regulates gene expression during zebrafish neurogenesis^38^, and ZNF460, which was pulled down only at the NPC stage and has largely unknown functions during neural development. For each protein, we established homozygous degron alleles in ESCs harboring HA-tagged endogenous EHMT1 to enable efficient EHMT1 pulldown in the absence or presence of a given candidate recruitment factor (**Figure 3C**). All degrons achieved full protein depletion after ESC culture with dTAG, as measured by a V5 tag included in our knock-in design (**Figure 3C,D, S3A**).

We conducted RNA-seq experiments with ZNF TF degron ESCs after 48 h of dTAG culture. Among the three candidates, ZNF462 loss most closely phenocopied EHMT1 depletion, whereas ZNF644 affected a smaller, distinct subset of EHMT1-responsive genes, and ZNF460 loss had little effect (**Figure 3E, S3B**). These observations were reflected in the overall impact of ZNF depletion on the ESC transcriptome: ZNF462 (858 DEGs) > ZNF644 (388 DEGs) > ZNF460 (156 DEGs) (padj <0.05, |log2fc|>0.5) (**Figure S3C**). The fact that depletion of each ZNF TF also affected genes that did not overlap with DEGs upon EHMT1 depletion (**Figure S3D**) documents partly EHMT1-independent functions of these transcriptional regulators.

ChIP-seq revealed that ZNF462 depletion reduced EHMT1 occupancy at nearly a quarter of EHMT1 peaks in ESCs (10,533/43,134), while ZNF644 depletion affected only 1,008 EHMT1 peaks (**Figure 3F,G**). These two largely non-overlapping sets of peaks were only bound by the corresponding ZNF TF (**Figure 3F, Supplementary Table 3**), suggesting independent roles for ZNF462 and ZNF644 in regulating EHMT1 genome occupancy. Similarly, genome-wide binding of these two ZNF TFs showed limited overlap (**Figure S3E**). We did not analyze the impact of ZNF460 depletion in this manner because of its limited impact on EHMT1 target genes (**Figure S3C**). In support of distinct regulatory functions of ZNF462 and ZNF644, EHMT1 peaks dependent on, and bound by, ZNF462 were strongly enriched for designated distal and intronic enhancers (**Figure 3H,I**), consistent with the presence of distinct H3K27ac peaks at these sites (**Figure 3J**). In contrast, EHMT1 peaks solely bound by ZNF644 were enriched for promoters (**Figure 3H,I**). Peaks bound by both ZNF462 and ZNF644 were also enhancer-enriched but did not lose EHMT1 upon depletion of either protein (**Figure 3G, S3F**), suggesting redundancy at those overlapping regions. Loss of either ZNF factor also increased EHMT1 occupancy at sites bound by the other, possibly reflecting redistribution of a limiting nuclear EHMT1 pool (**Figure 3G, S3F**).

Sites of ZNF462-dependent EHMT1 binding were significantly enriched near DEG^up^, indicating their regulatory significance (**Figure S3G,H**). This included several genes with established roles in neurodevelopment and/or NDD, which showed ZNF462-dependent EHMT1 binding at distal H3K27ac-marked candidate enhancer regions (**Figure 3J, S3I**). We also noted that EHMT1 peaks bound and/or regulated by ZNF462 or ZNF644 were among the strongest peaks based on ChIP-seq signal (**Figure S3J**). This might reflect efficient ZNF-dependent recruitment of EHMT1 to a comparably small subset of regulatory sites. In contrast, the more abundant weak, ZNF462/644-independent EHMT1 peaks might reflect a distinct molecular mechanism, such as spreading via EHMT1’s ability to bind its own H3K9me2 mark^39^, although we cannot rule out that another TF such as WIZ is active at these sites. Thus, ZNF462 and ZNF644 play distinct roles in EHMT1 retention at chromatin in self-renewing ESCs, with ZNF462 playing a major role at enhancer regions near neurodevelopmental gene loci, reminiscent of observations in mouse ESCs^35^. Overall, these findings establish ZNF462 and ZNF644 as distinct determinants of EHMT1 genomic targeting in ESCs and identify broadly expressed zinc finger proteins as candidate mediators of developmental rewiring of EHMT1 recruitment.

### ZNF462 drives developmental rewiring of EHMT1 genome occupancy during neural induction

So far, our findings document that ZNF462 contributes to EHMT1 retention at a subset of distal enhancers in ESCs. To query whether this transcription factor also drives developmental remodeling of the EHMT1 cistrome during exit from pluripotency and neural induction, we differentiated EHMT1-dTAG and ZNF462-dTAG ESCs for three days in dual SMAD and WNT inhibition (“d3 NE”)(**Figure S4A**), followed by HA (EHMT1) and V5 (ZNF462) ChIP-seq. K-means clustering (**Figure 4A**) revealed 4,055 EHMT1 peaks that strongly gained (“d3-specific”; cluster C3) and 2,259 strong peaks that lost (“ES-specific”; cluster C4) signal in early NE. The majority of these peaks – as well as a larger number of regions that showed similar trends, albeit less pronounced (C1 and C2, respectively; **Figure 4A**) – localized to distal regions, while a smaller group of mostly unchanged peaks (C5) associated with promoters (**Figure S4B**). This clustering demonstrates a substantial degree of regulatory reprogramming among EHMT1 target sites during early neural induction, which is accompanied by largely distinct transcriptional responses to EHMT1 depletion in d3 NE and ESCs (**Figure S4C**).

**Figure 4.**
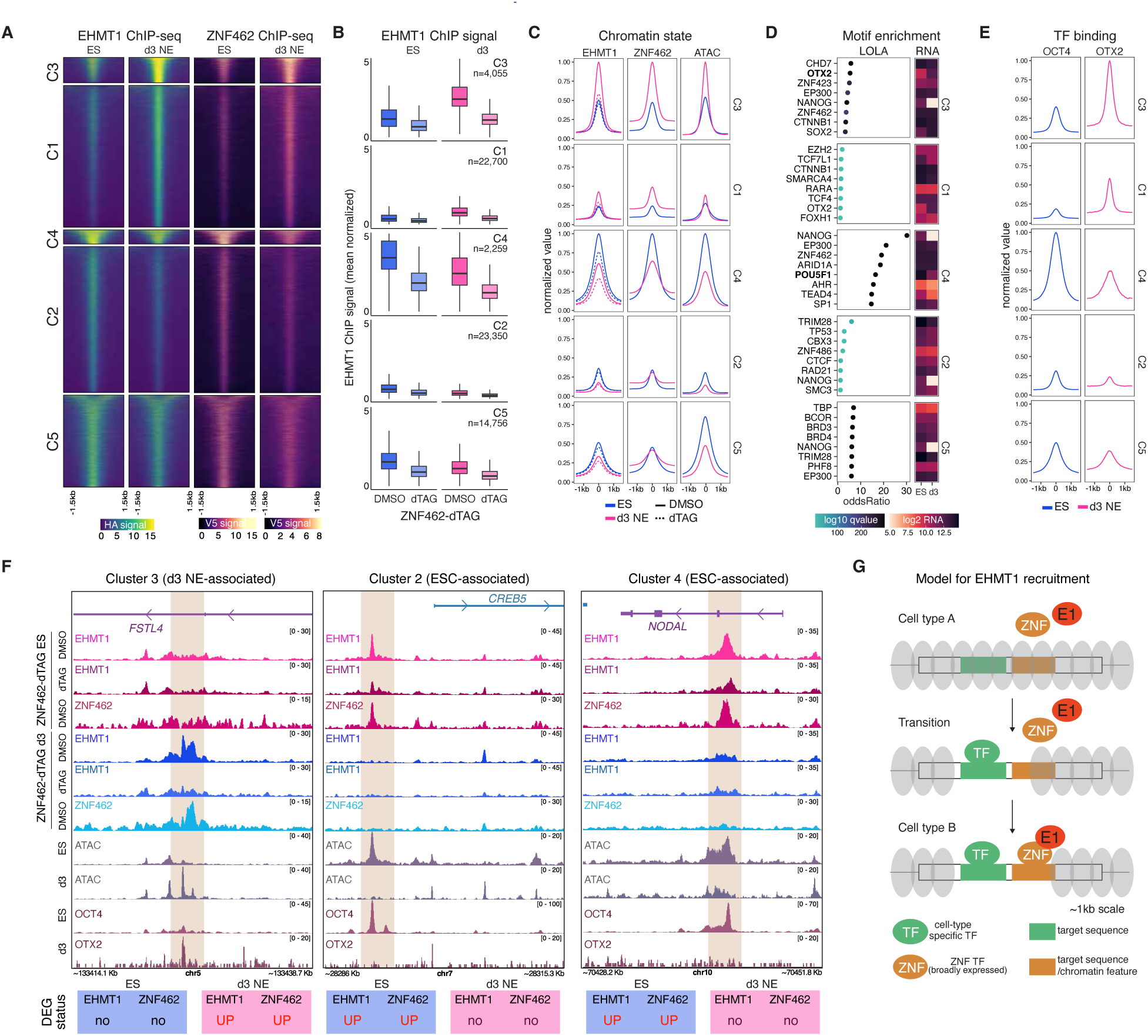
ZNF462 drives developmental rewiring of EHMT1 genome occupancy during neural induction. **A**, K-means clustered heatmap of EHMT1 and ZNF462 ChIP signal in ES and day-3 neuroectoderm (d3 NE). **B,** Box plot of EHMT1 ChIP signal for each k-means cluster under ZNF462 depletion in ES and d3 NE. Wilcox test p-value <2.2e-16 for all DMSO-dTAG comparisons. **C,** Average profile plots of normalized EHMT1, ZNF462, and chromatin accessibility (ATAC) signal for each k-means cluster in ESCs (GSE109524) and d3 NE. Dashed lines represent signal following ZNF462 depletion. **D,** LOLA motif analysis using all EHMT1 peaks in ES and d3 NE as the background set. The heatmap shows expression levels (log2-normalized RNA-seq counts) of enriched factors in ES and d3 NE. **E,** Average profile plots of OCT4 ChIP signal (GSM3381968) in ES and OTX2 CUT&RUN signal in d3 NE. **F,** Representative genome tracks showing key chromatin features at different EHMT1 peak categories (orange box highlight). EHMT1-dTAG and ZNF462-dTAG DEG status are shown below (DESeq2 padj < 0.05, |log2FoldChange| > 0.5). **G,** Model of cell type-specific EHMT1 recruitment by ZNF TFs and stage-specific transcription factors.

ZNF462 genome occupancy changes during the ES-to-NE transition closely mirrored those observed for EHMT1 (**Figure 4A**), suggesting coordinated redistribution of these two regulators. Moreover, ZNF462 depletion strongly impaired the establishment of C3 EHMT1 peaks in d3 NE while accelerating the loss of ES-specific (C4) peaks (**Figure 4B,C, S4D**), supporting a central role of ZNF462 in driving the observed remodeling of EHMT1 genome occupancy. Since *ZNF462* mRNA levels were insensitive to the signaling cues driving anterior neural induction (**Figure S4E**), stage-specific redistribution is unlikely to be primarily driven by changes in its abundance. Instead, the close correlation between ZNF462 and EHMT1 genome binding and dynamic chromatin accessibility (**Figure 4C, S4D**) suggested that changing lineage-specific chromatin landscapes might redirect ZNF462 binding. Indeed, LOLA ChIP enrichment of the identified binding clusters revealed lineage-associated TFs matching the stage-specific cistrome changes. OTX2, an essential regulator of forebrain and midbrain development^40,41^, was among the top-enriched motifs for the d3-specific cluster C3 (**Figure 4D**) and was strongly upregulated during neural induction (**Figure S4E**). In contrast, the pluripotency-associated pioneer factor POU5F1 (OCT4)^42^ was enriched at ES-specific C4 peaks (**Figure 4D**) and experienced sharp downregulation during differentiation (**Figure S4E**). Importantly, CUT&RUN for OTX2 showed strong binding of this TF in d3 NE at C3 peaks (**Figure 4E, S4D**), while analysis of public ChIP-seq data^43^ confirmed the presence of OCT4 at C4 peaks in ES.

The correlation between stage-specific binding of transcriptional regulators, chromatin accessibility, and transcriptional response to ZNF462 and EHMT1 loss was evident at gene loci encoding several regulators of epiblast differentiation and neuronal development, including the BDNF repressor FSTL4 operational at later stages of neural development^44^ (an EHMT1/ZNF462 target only in d3 NE), the neural stem cell regulator CREB5^45^ (a target only in ESC), and the neural induction antagonist NODAL^46^ (a target only in ESC) (**Figure 4F**). Together with the absence of lineage-specific TFs among EHMT1 interaction partners (**Figure 3A**), these observations support a model for cell type-specific EHMT1 target recognition in which opportunistic binding of ZNF TFs within changing chromatin landscapes associated with signaling-controlled, lineage-associated transcription factors mediates remodeling of EHMT1 to stage-specific regulatory elements (**Figure 4G**).

### ZNF462-dependent EHMT1 recruitment shapes shared and distinct transcriptional programs during NPC specification

Having established that ZNF462 contributes to the developmental remodeling of EHMT1 genome occupancy during human neural induction, we next asked how disrupting this recruitment mechanism affects transcriptional programs during later stages of cortical development. Specifically, we examined how prolonged deficiency of either EHMT1 or ZNF462 affects global gene expression in NPCs, a critical intermediate in cortical neurogenesis, which we found to be largely refractory to short-term EHMT1 loss (see **Figure 1E**). Of note, EHMT1 or ZNF462 depletion from the ESC to the d20 NPC stage (**Figure S5A,B**) yielded morphologically indistinguishable DMSO and dTAG cultures with similarly high percentages of PAX6^+^ cells (**Figure S5C,D** and data not shown), indicating that human pluripotent cells remain capable of differentiating into NPCs despite drastic reductions in EHMT1 or ZNF462 levels.

RNA-seq after prolonged depletion of either EHMT1 or ZNF462 identified several hundred significantly dysregulated genes (|log2FC|>0.5, padj <0.05) in NPCs, with upregulated genes predominant for either protein (509/773 for EHMT1, 312/500 for ZNF462) (**Figure S5E**). Although direct overlap between EHMT1 and ZNF462 DEGs was modest (**Figure S5F**), clustering of all affected genes yielded a more nuanced picture and revealed a strongly co-dysregulated subset of transcripts. This cluster (G5) comprised a mix of genes associated with either neuronal function – involving both excitatory and inhibitory neurotransmission and including NDD-associated genes such as IGF1^47^, DRD2^48^, and GLRA2^49^ – or the response to cellular stresses, including interferon-induced genes such as OAS3, BST2, IFITM1, and IFITM2^50,51^ (**Figure 5A-C**). Several genes and programs that were dysregulated after prolonged depletion already showed concordant but weaker changes in d3 NE and, to a lesser extent, ESC (**Figure 5C**). This supports the idea that while EHMT1 functions are primarily stage-specific, molecular abnormalities initiated at earlier stages can be amplified during continued development.

**Figure 5.**
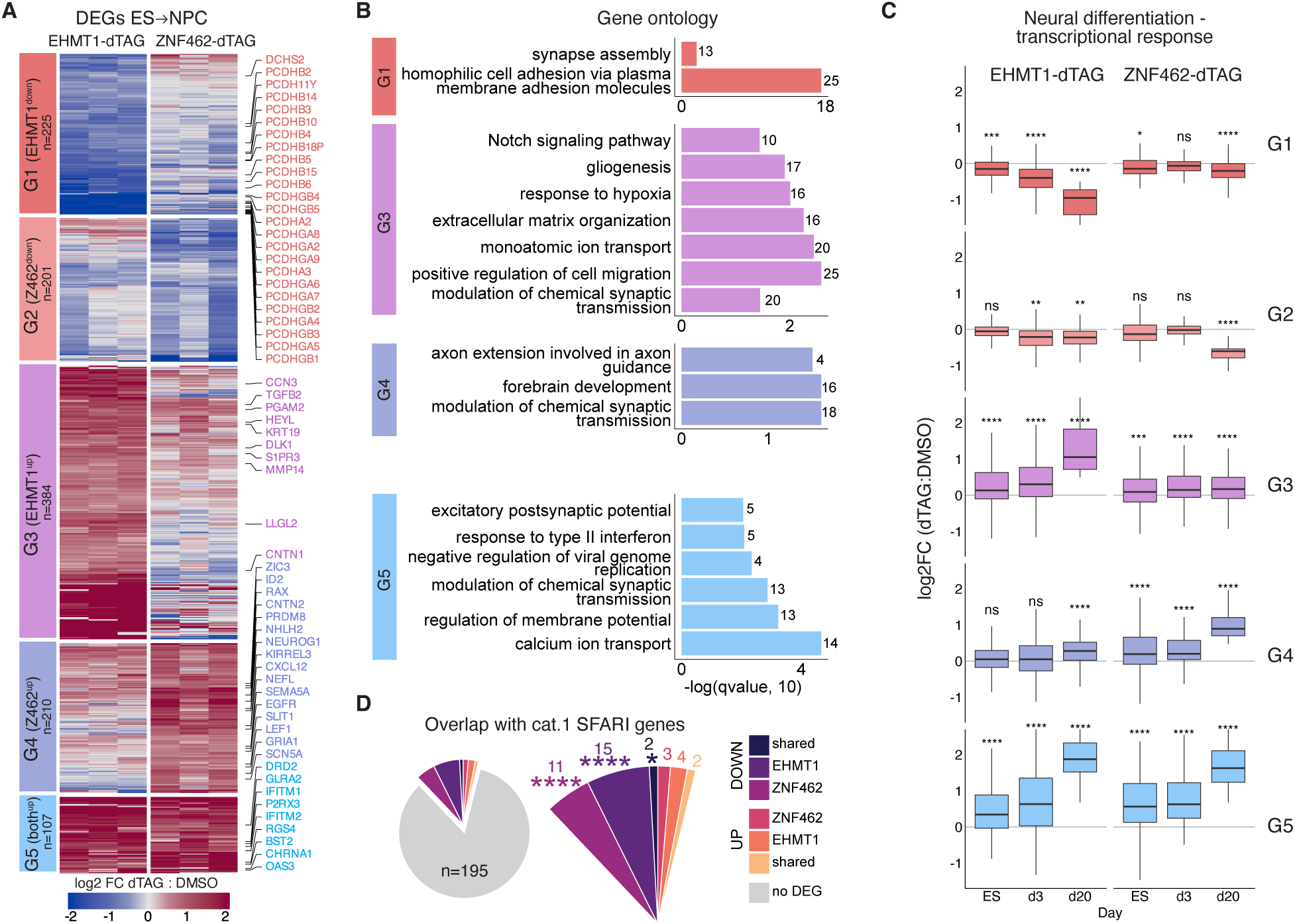
ZNF462-dependent recruitment underlies shared and distinct EHMT1 transcriptional programs during NPC specification. **A**, K-means heatmap showing RNA-seq expression changes at EHMT1 and ZNF462 DEGs (padj < 0.05, |log2FoldChange| > 0.5) upon depletion from the ES to d20 NPC stage. Genes assigned to select GO terms are highlighted: G1, homophilic cell adhesion via plasma membrane adhesion molecules (GO:0007156); G3, Notch signaling pathway (GO:0007219); G4, forebrain development (GO:0030900); G5, excitatory postsynaptic potential (GO:0060079) and negative regulation of viral genome replication (GO:0045071). **B,** GO enrichment terms for each k-means cluster of DEGs. The numbers of DEGs within each term are indicated. **C,** RNA-seq expression changes of the genes in the indicated k-means clusters after EHMT1 or ZNF462 depletion at the ES stage, from ES to d3 NE, and from ES to d20 NPC; one-sample t-test with tdf=n-1, 95% CI, Cohen’s D against the log2FC = 0 null hypothesis, corrected using Bonferroni. **D,** Pie chart showing overlap between Category 1 SFARI genes and genes affected by prolonged (ES to NPC) EHMT1 or ZNF462 depletion (padj < 0.05, |log2FoldChange| > 0), hypergeometric test (see methods for details).

Beyond this shared core, EHMT1 and ZNF462 deficiency affected largely distinct developmental programs. For example, NPCs lacking EHMT1 upregulated cell migration regulators (G3) and – consistent with observations made in mouse models – downregulated neurodevelopmentally important cell adhesion molecules such as proto-cadherins^52^ (G1), while NPCs lacking ZNF462 showed evidence for dysregulated axial patterning (G4) (**Figure 5B**). EHMT1– or ZNF462-repressed genes (G3, G4) also showed evidence of functional convergence, including modulation of synaptic transmission (**Figure 5B**). Genes uniquely affected by EHMT1 or ZNF462 depletion, and to a lesser extent shared targets, were significantly enriched for ASD-associated genes relative to the NPC transcriptome (**Figure 5D, S5G**).

Together, these observations show that EHMT1 and ZNF462, despite their close biochemical association within the same protein complex and shared genomic targets during neural induction, control only partially overlapping transcriptional programs during NPC specification. These findings support partial convergence between EHMT1– and ZNF462-dependent regulatory networks while emphasizing factor-specific functions. These patterns of molecular convergence and divergence are consistent with the overlapping neurological manifestations of the Weiss-Kruszka (ZNF462) and Kleefstra (EHMT1) syndromes, including developmental delay and autistic features, as well as their distinct clinical features^4,36,53,54^.

### Developmental EHMT1 deficiency disrupts neuronal maturation and transcription in a subtype-specific manner

Having found evidence for a partially cumulative effect of EHMT1 loss throughout NPC specification, we sought to determine how these developmental perturbations manifest in neurons. To address this question, we performed transcriptional profiling of d35 cultures exposed to dTAG from the ES stage onward (**Figure S6A**), applying single-cell RNA-seq (scRNA-seq) to capture heterogeneity introduced by prolonged culture.

As expected, d35 cells were composed predominantly of postmitotic neurons expressing *MAP2*, *DCX*, *and VAMP2*, with elevated *FOXG1 levels* consistent with a telencephalic origin, and lacking proliferative and progenitor markers (**Figure S6B**). In addition to *SLC17A6*^+^ excitatory neurons^27^, the cultures contained substantial numbers of *GAD2*^+^ inhibitory neurons and rare *NFIA^+^* glioblasts (**Figure 6A, S6B,C**). Marker-based annotation resolved distinct neuronal populations, including *FOXP2*^+^ excitatory cortical projection neuron-like cells (hereafter CPN)^55^, *NR2F2*^+^ CPN2^56^, and several inhibitory populations with medium spiny neuron-like identities, including *EBF1*^+^ D1-MSNs, *SIX3*^+^ D2-MSNs, and *ISL1*^+^ D1/D2-MSNs^57,58^, as well as *NDNF*^+^ layer 1 interneuron-like cells (hereafter L1 INT)^59^ (**Figure 6A,B**, **S6D**). These observations allow us to compare the impact of EHMT1 depletion across multiple transcriptionally distinct neuronal populations.

**Figure 6.**
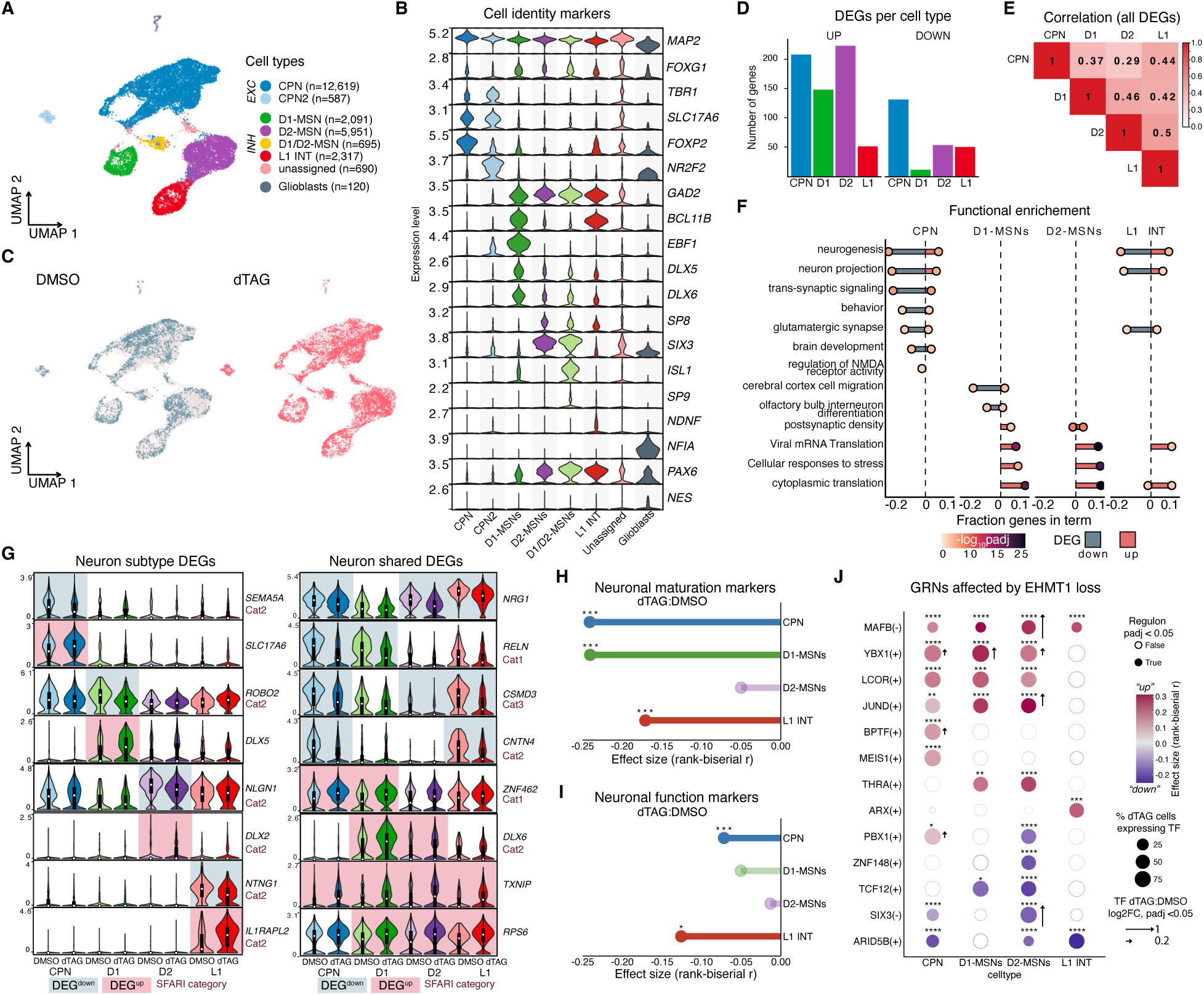
Developmental EHMT1 deficiency affects cortical neurons in a subtype-specific manner. **A**, scRNA-seq UMAP of d35 CN colored by assigned cell type, excitatory (EXC) and inhibitory (INH) neuronal subtypes, glioblasts, and unassigned cells. The numbers of cells per cluster are shown in parentheses. **B,** Violin plots of Seurat-normalized expression levels for cell-identity marker genes. **C,** UMAPs of d35 CNs exposed to different treatments (DMSO = blue; dTAG = red). **D,** DEGs per cell type after ES→CN depletion, as identified with Seurat FindMarkers (padj < 0.05, |log2FoldChange| > 0.5). **E,** Degree of similarity of combined DEGs identified in the indicated neuronal cell types, as indicated by log2FoldChange pairwise Pearson correlation. **F,** Functional enrichment (gProfiler; padj < 0.05) for ranked DEGs in the indicated neuronal subtypes. **G,** Violin plots showing subtype-specific (left) and shared (right) DEGs between neuronal subtypes. Upregulated DEGs highlighted in red and downregulated DEGs highlighted in blue. **H,** Impact of EHMT1 depletion on the neuronal maturation score in neuronal subtypes (see Methods). Mann-Whitney test with Bonferroni correction. **I,** As H for neuronal function score (see Methods). **J,** Transcription factors with regulons significantly affected by EHMT1 depletion (Mann-Whitney test with Bonferroni correction), as determined by SCENIC (see Methods). (+)(-) indicate positive and negative correlations between the expression of a transcription factor and its regulon, respectively. Arrows indicate significant (padj<0.05) log2FC (dTAG/DMSO) of the TF in the cell type.

All major neuronal populations were recovered in ES→CN depletion cultures, but DMSO and dTAG cells showed distinct distributions within the UMAP cell populations (**Figure 6C**), suggesting deregulated neuronal programs in the absence of EHMT1. We restricted differential gene expression analysis to CPN, D1-MSN, D2-MSN, and L1 INT, all of which were represented by at least 2,000 cells (**Figure 6D**). At the level of individual genes, EHMT1 deficiency produced largely distinct transcriptional responses across these four cell populations (**Figure S6E, Supplementary Table 4**) with low correlation scores (**Figure 6E**). Nevertheless, functional enrichment analysis revealed some degree of convergence in broader cellular programs. In CPN, EHMT1 deficiency predominantly reduced programs associated with neurogenesis, trans-synaptic signaling, and neuron projection, with some aspects of these changes, including specific targets, shared with L1 INT (**Figure 6F, S6F**). In contrast, inhibitory neuronal subtypes predominantly showed the induction of cytoplasmic translation and viral mRNA translation programs **(Figure 6F, S6F**), consistent with activation of stress-associated transcriptional programs by long-term EHMT1 depletion.

The subtype-specific responses to EHMT1 deficiency involved genes with established roles in neurodevelopment or neuronal function, including dysregulation of the autism-associated *RELN*^60^ and *ZNF462* in both CPN and D1-MSNs (**Figure 6G, S6G**), upregulation of *SLC17A6*, which encodes a facilitator of glutamate secretion, in CPN, and of several members of the DLX family of lineage specification and differentiation regulators in D1 and D2 MSNs. Consistent with these gene-level changes, established cortical neuron maturation and neuronal functionality modules^27^ were differentially perturbed across neuronal subtypes (**Figure 6H,I**), supporting the notion that EHMT1 influences the timing of human cortical neurogenesis by regulating subtype-specific transcriptional programs.

As our earlier analyses had indicated that EHMT1 is integrated into cell-type-specific TF networks, we asked whether prolonged EHMT1 deficiency alters distinct regulatory networks across neuronal cell types. SCENIC analysis, which infers both activating and repressing modules from scRNA-seq^61^, revealed both shared and subtype-specific changes in regulon activity (**Figure 6J**). For example, genes in a regulon repressed by the brain development regulator MAFB^62,63^ and genes in regulons activated by JUND and YBX1, two developmental TFs also involved in adaptive cellular stress responses^64,65^, showed increased expression across multiple neuronal cell types (**Figure 6J**). Among cell type-specific responses, we observed altered output of regulons associated with neurogenesis-associated TFs such as ARX^66^ in L1 interneurons, THRA^67^ in D1/D2-MSNs, and PBX1^68^ in D2-MSNs (**Figure 6J**). Notably, only a subset of gene loci encoding the corresponding TFs, including *YBX1*, changed in expression upon EHMT1 depletion (**Figure 6J**), indicating that altered regulon output was not generally caused by altered TF abundance. Thus, EHMT1 function is coordinated with multiple transcription factor networks during later stages of neurogenesis, providing a regulatory context for both shared and distinct transcriptional consequences of prolonged EHMT1 deficiency across neuronal subtypes.

### Developmental EHMT1 deficiency causes cumulative but largely reversible transcriptional dysregulation

The gene expression abnormalities observed during prolonged (ES-to-d35) EHMT1 deficiency prompted us to determine when these molecular alterations arise during differentiation. Examination of developmental gene expression trajectories revealed that d35 DEGs upregulated in inhibitory cell types (D1 MSN, D2 MSN, L1 INT) were expressed at high levels throughout differentiation, whereas genes upregulated in CPNs showed lower expression levels that peaked in NPCs (**Figure 7A**). In contrast, all four sets of genes downregulated in the major neuronal cell types present in d35 cultures showed marked developmental upregulation from the NPC stage onward (**Figure 7A**), suggesting that NPC specification may represent the developmental window during which EHMT1-dependent neuronal programs are established. We therefore extended our scRNA-seq experiments to include cells that only experienced EHMT1 depletion starting at the NPC stage (“NPC→CN”) (**Figure 7B**).

**Figure 7.**
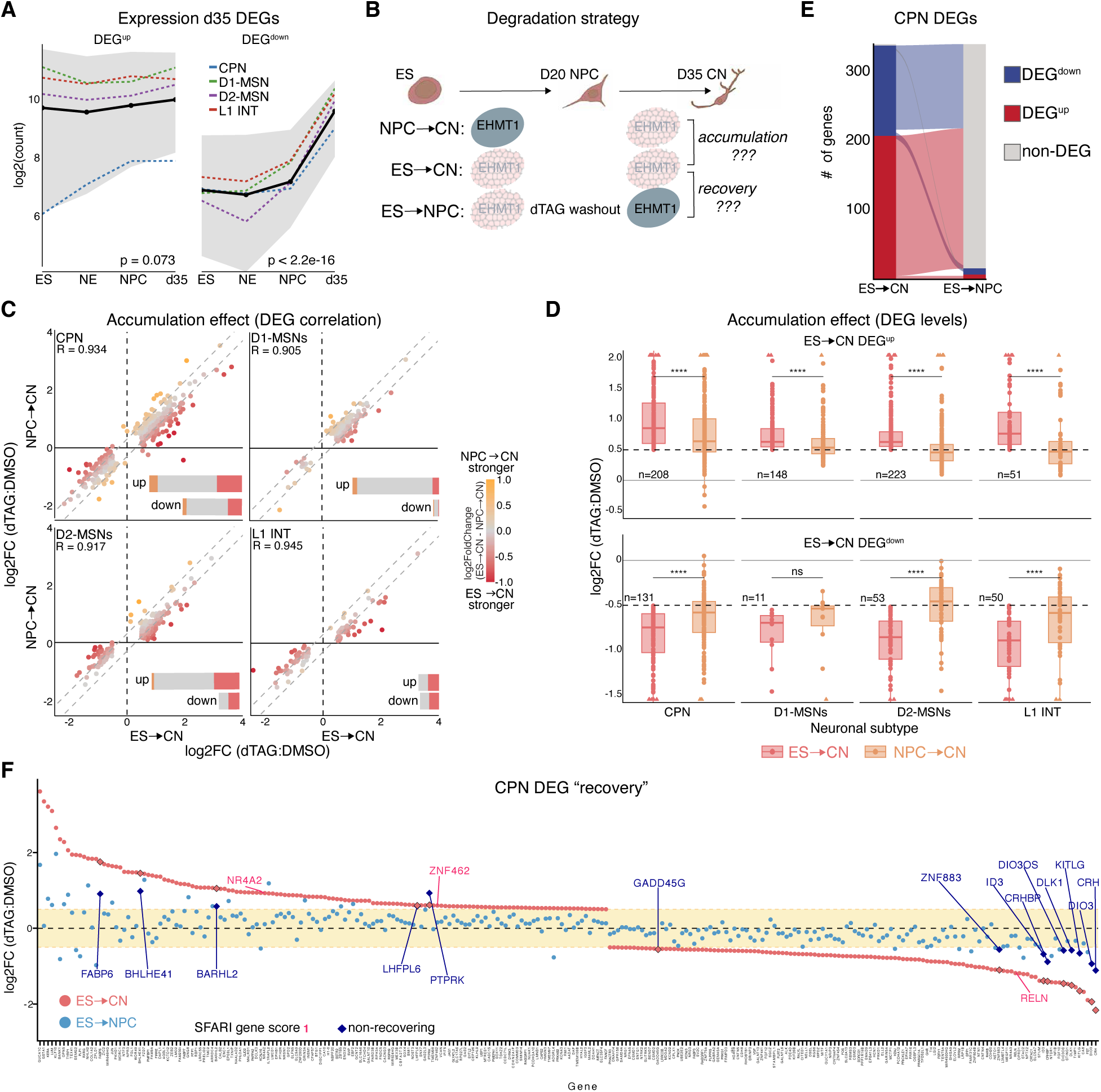
Accumulation and reversibility of transcriptional dysregulation caused by developmental deficiency of EHMT1. **A**, Expression trajectories of ES◊CN DEGs (padj <0.05, |log2FoldChange| > 0.5) identified in different neuronal subtypes (CPN, D1-MSN, D2-MSN, L1 INT; dotted line) across neural development. The black line shows the mean across subtypes and the shaded area the quartile range. Significance tested for variation between mean counts across stages – Kruskal-Wallis p-values indicated. **B,** Experimental strategy to compare EHMT1 deficiency throughout neural development (ES◊CN) to EHMT1 deficiency until the NPC stage (ES◊NPC) or from the NPC stage onwards (NPC◊CN). **C,** Correlation of gene expression changes triggered by ES◊CN and NPC◊CN EHMT1 depletion at EHMT1-controlled genes in different neurons. Pearson R value shown per neuronal subtype. Bar plots indicate the number of DEGs with |log2FoldChange(ES◊CN – NPC◊CN)| >0.3 (red), <-0.3(orange) and –0.3-0.3 (grey). **D,** Comparison of the RNA expression changes of all ES◊CN DEGs after ES◊CN and NPC◊CN depletion in the different cell types. Triangles at the plot edge indicate outliers; paired t-test. **E,** Alluvial plot of ES◊CN and ES◊NPC DEGs (padj <0.05, |log2FoldChange| > 0.5) in CPNs. Color indicates DEG status (red=up, blue=down, grey=non-DEG). **F,** Dot plot comparing RNA expression level changes of ES◊CN DEGs in CPNs after EHMT1 depletion from ES◊CN to ES◊NPC. Genes that remain significantly affected (padj <0.05, |log2FoldChange| > 0.5) upon partial EHMT1 recovery are highlighted with dark blue diamonds. Category1 SFARI genes are highlighted in red.

UMAP densities suggested a high degree of similarity between ESC→CN– and NPC→CN-depleted cells compared to the DMSO control (**Figure S7A**), and, more importantly, the transcriptional responses were highly correlated across all four neuronal subtypes (R > 0.9). This indicates that EHMT1 loss beginning at the NPC stage largely perturbed the same programs as depletion from the ESC stage onwards (**Figure 7C**). However, the magnitude of transcriptional dysregulation (**Figure 7D**) and the number of called DEGs (**Figure S7B**) were consistently greater after ESC→CN depletion. In addition, GO analysis (**Figure S7C**) and maturation/function scores (**Figure S7D**) indicated a qualitatively similar but stronger effect of ESC→CN depletion on transcriptional programs regulated by EHMT1. Together, these observations document a cumulative quantitative effect of prolonged EHMT1 deficiency on gene expression during neurogenesis, manifesting as the magnitude of observed dysregulation rather than substantial differences in the neuronal programs affected.

Having established that EHMT1 deficiency beginning at the NPC stage is sufficient to disrupt many of the same neuronal programs as continuous depletion from ESCs onwards, we next asked whether restoring EHMT1 protein at this stage can rescue transcriptional abnormalities caused by earlier developmental deficiency. To test this, we restored EHMT1 beginning at the NPC stage by washing out dTAG (**Figure 7B**), resulting in a recovery of about 70% of EHMT1 levels by d35 (**Figure S7E,F**). Surprisingly, UMAP density comparison revealed that ‘washout’ CNs (ESC→NPC) localized together with cells that had never experienced EHMT1 depletion, suggesting widespread resetting of their transcriptome (**Figure S7G**). In addition, comparison of called DEGs (padj <0.05, |log2fc| >0.5) revealed nearly complete absence of statistically significant transcriptional dysregulation at the individual gene level in all neuronal subtypes (**Figure 7E, S7H-J**). Projection of the washout-to-DMSO expression ratio for genes dysregulated after continuous ESC→CN depletion showed widespread movement back toward control expression following EHMT1 restoration, including at most NDD-associated gene loci (**Figure 7F, S7K-N**). The few genes with incomplete recovery generally were more strongly dysregulated by EHMT1 depletion (**Figure S7O**), suggesting elevated sensitivity to EHMT1 dosage, although we cannot rule out the introduction of lasting epigenetic alterations at these loci. We confirmed these recovery effects in independent bulk RNA-seq experiments (**Figure S7P,Q**). Thus, EHMT1 retains the ability to engage effectively with and appropriately regulate most genes and stage-specific programs dysregulated by prolonged developmental deficiency, revealing substantial transcriptional plasticity.

## Discussion

Our findings show that the essential histone methyltransferase EHMT1 does not perform a fixed regulatory function during human cortical development but is repeatedly repurposed through stage-specific changes in genomic targeting and transcriptional output. More broadly, these observations suggest that developmental progression fundamentally reshapes the regulatory logic by which a ubiquitously expressed chromatin regulator influences gene expression, thereby allowing the same protein to fulfill distinct functions across successive developmental contexts. Together with studies in mouse ESCs^35^, our short-term (48 h) depletion experiments support a conserved role of EHMT1 in suppressing developmental gene expression programs, including those associated with germ-layer differentiation and neurogenesis, in the pluripotent state. However, during neural induction and thereafter, EHMT1 also supports the expression of neuronal function-associated programs across multiple neuronal subtypes. Whereas transient enzymatic inhibition of EHMT1/2 in cortical NPCs was previously shown to accelerate aspects of subsequent neuronal maturation^27^, our temporally resolved EHMT1 depletion experiments reveal that prolonged EHMT1 deficiency ultimately compromises faithful neuronal maturation. Together, these findings underscore the developmental context dependence of EHMT1 function and support altered cellular maturation as a phenotypic convergence point across distinct ASD-associated proteins^11^. In inhibitory neurons in particular, these maturation defects are accompanied by activation of cellular stress-response programs, as previously indicated for different cell types in EHMT1 heterozygous mouse brains^69^. Thus, although EHMT1 regulates distinct sets of genes at successive developmental stages, its prolonged absence during neural differentiation ultimately disrupts broader, partially shared programs in neuronal subtypes.

A second major finding is that EHMT1 occupancy alone poorly predicts transcriptional sensitivity to its loss. At the cellular level, the greater number of dysregulated genes in ESCs than in NE and NPCs upon acute EHMT1 depletion may reflect progressive stabilization of H3K9me2-marked chromatin as cells differentiate and proliferative chromatin dynamics decline^70^. By contrast, the comparatively strong acute response in postmitotic neurons suggests that EHMT1-dependent regulation at this stage may involve more dynamic maintenance of histone marks^71^, altered cofactor requirements, or functions not explained solely by replication-coupled H3K9me2 turnover. At the genomic level, EHMT1 occupies substantially more regulatory elements, including enhancers and promoters, than show transcriptional responses to EHMT1 loss. This discrepancy persists even within broader H3K9me2 domains. Thus, sensitivity to EHMT1 loss appears to depend on both localization within chromatin environments permissive to EHMT1 binding and the stable propagation of elevated H3K9me2 levels, as well as a transcriptionally quiescent state rendered competent by the availability of appropriate trans-acting factors. Consequently, EHMT1 fine-tunes the activity of developmental gene loci in a stage-dependent manner rather than acting as a binary on/off switch. This model is consistent with observations in other model systems following loss of EHMT1/2 activity^35^, as well as with our identification of cell type-specific gene regulatory networks whose activity changes following EHMT1 loss.

Our findings further suggest that developmental repurposing of EHMT1 is not primarily achieved by replacing its core chromatin-associated machinery. Instead, EHMT1 exhibits remarkable flexibility in enhancer occupancy and target gene regulation during human cortical differentiation, despite a relatively stable set of core protein partners across cell types. We propose that EHMT1 achieves this flexibility through interactions with a limited number of broadly expressed zinc finger proteins that act within chromatin landscapes pioneered by lineage-specific transcription factors^72^. This model provides a potentially generalizable mechanism by which ubiquitously expressed chromatin modifiers can acquire developmental specificity through broadly expressed recruitment factors operating within changing lineage-specific regulatory landscapes. It remains to be determined whether ZNF462 and related factors recruit EHMT1 through direct recognition of specific DNA motifs, through interactions with other chromatin-bound proteins, or through features of the local chromatin environment. Consistent with studies in mouse ESCs^31,35^, our findings support a central role for ZNF462 in recruiting EHMT1 to developmental enhancers, whereas ZNF644 acts at promoters in human ESCs. ZNF462 and ZNF644 have also been found to be associated with EHMT1/2 in human kidney cells and the fish retina^38^. The use of related zinc finger proteins to recruit the EHMT1/2 complex across different tissues and species suggests that this recruitment logic may represent a broadly conserved mechanism. Further studies are needed to elucidate the mechanism and factors involved in EHMT1 recruitment to the many genomic sites unaffected by loss of ZNF462 or ZNF644.

Our findings also might have implications for understanding neurodevelopmental disorders caused by chromatin-regulator haploinsufficiency. The overlap between EHMT1– and ZNF462-controlled transcriptional programs in NPCs supports a partially convergent neurodevelopmental regulatory network but also shows that loss of even functionally closely related proteins can cause distinct molecular consequences within the same developmental cell state. Strikingly, although we observe greater gene dysregulation in neurons upon EHMT1 loss from the epiblast than from the NPC stage onward, almost all loci dysregulated after prolonged EHMT1 deficiency shifted back toward control expression levels after restoration of near-physiological EHMT1 levels. This documents that EHMT1 remains capable of re-engaging stage-appropriate targets after prolonged deficiency. Although additional studies are needed to determine whether transcriptional and epigenetic recovery are complete, these findings indicate that many neuronal transcriptional programs remain molecularly recoverable despite extended developmental disruption. This degree of molecular reversibility is consistent with studies in mouse models of Kleefstra syndrome^69^ and MECP2 deficiency^73,74^, which show that selected neurological phenotypes can be ameliorated by subsequent restoration of gene function. The small number of refractory loci we identified may provide a tractable model for identifying the chromatin features underlying transcriptional scarring. Together, our findings establish developmental rewiring of recruitment and transcriptional output as a mechanism by which ubiquitously expressed chromatin regulators can acquire stage-specific functions and reveal that developmental gene regulatory programs can retain substantial capacity for molecular recovery following prolonged loss of a disease-associated regulator.

## Resource Availability

### Lead contact

Requests for further information and resources should be directed to and will be fulfilled by the lead contact, Matthias Stadtfeld.

## Materials availability

All unique/stable reagents generated in this study are available from the lead contact upon completion of a materials transfer agreement.

## Data and code availability

Genomics and epigenomic data have been deposited at GEO for RNA-seq, ChIP-seq, and single-cell RNA-seq under accession numbers GSEXXXXX, GSEXXXXX, and GSEXXXXX, respectively. The data are currently private and will be made publicly available upon formal journal publication. Pipelines and scripts used for analysis and figure generation in this paper are available through Zenodo at https://doi.org/10.5281/zenodo.21925204. All datasets and code will be publicly available as of the date of publication, and reviewer tokens have been provided. Proteomics data have been deposited in the ProteomeXchange Consortium via the PRIDE partner repository with the dataset identifier PXD082029.

## Acknowledgements

We thank Oliver Bell, Gabriele Ciceri, Nan Xu, Uk Jin Lee, and all members of the Stadtfeld and Apostolou labs for their valuable input to this study. We thank the Weill Cornell Medicine Genomics Resources Core Facilities for expert assistance. The Sidoli lab gratefully acknowledges funding from the NCI (1R21CA316284), the Aging Biology Foundation, the Hevolution Foundation (AFAR), the ERCM-CFAR Center for AIDS Research, and the NIH Office of the Director (S10OD030286). E.A. was supported by R01NS136475. L.S. was supported in part by grants R01MH135403 and R01NS128087 and by core grant support from P30CA08748. M.S. was supported by grants from the NIH (R01MH139528), the Simons Foundation, the Tri-Institutional Stem Cell Initiative, and the Bohmfalk Charitable Trust.

## Author Contributions

K.K. established all degron lines, planned, executed, and analyzed all differentiation, bulk RNA-seq, H3K9me2 ULI-nChIP experiments, as well as HA and V5 ChIP-seq in ESCs. K.C. conducted HA and V5 ChIP-seq in d3 NE, V.B. helped with cortical cell type assignment and conducted scRNA-seq analysis, S.R.M., C.U., and A.P. provided genomics pipelines and advised on genomics analyses, W.M. conducted ATAC-seq in d3 NE, E.G. did OTX2 ChIP-seq in d3 NE, A.L.D. helped with histone proteomics, G.T. did cloning for ZNF degron lines, A.L.D, E.G., and W.M. helped with cell culture and gene targeting, A.M. helped with cortical differentiation and gene targeting, R.W. assisted in setting up human ESC targeting and cortical differentiation, E.A. helped plan and guide genomics and epigenomics experiments, S.S. conducted mass-spec experiments, L.S. provided guidance on execution and interpretation of differentiation experiments, M.S. conceived the study, provided funding, supervised experiments and analyzed data. K.K. and M.S. wrote the manuscript with input from all authors.

## Declaration of Generative AI

During preparation of this manuscript, the authors used Claude and ChatGPT to identify grammatical and typographical errors and improve the flow of the text. After using these tools, the authors reviewed and edited the content as needed and take full responsibility for the published article.

## Methods

### Human ESC culture

Experiments with human ESCs were approved in compliance with the Tri-Institutional ESCRO at Weill Cornell Medicine, Memorial Sloan Kettering Cancer Center, and Rockefeller University. MEL-1 ESCs were obtained from WiCell Stemcell Bank. Human MEL-1 ESCs were cultured on plates coated with vitronectin (Thermo Fisher Scientific (TF), A14700) in Essential 8 Flex media with supplement (TF A2858501) and normocin (50 ug/ml) (InvivoGen, ant-nr-05). Cells were passaged with home-made EDTA dissociation solution (0.5 mM EDTA and 35 mM NaCl in PBS1x) during maintenance or with Accutase (TF 00-4555-56) for freezing or plating for experiments. Cell stocks were frozen in Stem Cell Banker (Amsbio, 11890). After thawing and single-cell seeding, ROCK inhibitor (Y-27632) (Tocris Bioscience, 27632) was added to the cell culture at 10 μM for 18-24 h.

### Human cortical neuronal differentiation

ESCs were differentiated using a previously established protocol^27^. ESCs were plated at a density of 3×10^5^ cells/cm^2^ in Essential 8 flex media with 10 uM Y-27632 on Matrigel (354234, Corning) coated plates (d-1). On days 0-9 the culture media was replaced daily with Essential 6 medium (E6; TF A1516401) supplemented with 10 μM SB431542 (Tocris, 1614) and 100 nM LDN193189 (Stem Cell Technologies, 72142); with an additional 2 μM XAV939 (Tocris, 374) from days 0-2 to induce anterior neuroectodermal patterning. Between days 10-20, the culture media were replaced daily with N2/B27 medium comprising 1:1 Neurobasal (NB; TF 21103049):DMEM/F12(TF 11330057) basal medium supplemented with 1× N2 (TF 17502048) and B27 minus vitamin A (TF 12587010) to generate a neurogenic population of cortical NPCs. Plates for cortical differentiation were prepared starting at d17 with 15 ug/ul poly-ornithine (Sigma, P3655) diluted in PBS; coated overnight at 37C, which was washed twice with PBS at d18 before adding a 1:1 ratio of 4 ug/ul Laminin (Sigma 3400-010-02) and 4 ug/ul Fibronectin (R&D Systems 1918-FN) in PBS for 48 h at 27C. At d20 the cells were dissociated with Accutase for 30-45min at 37C and either frozen in Stem Cell Banker or plated to induce synchronized neurogenesis as follows: cells were counted and plated at 1.5×10^5^ cells/cm^2^ onto PO/L/F plates in NB/B27media (NB/B27-vitA + 1% L-glutamine + 1% penicillin-streptomycin) with 10uM Notch inhibitor (R&D 2634) added. We changed the medium every 4 days. From d30-35, media were supplemented with 20 ng/ml BDNF (PeproTech 450-10), 20 ng/ml GDNF (R&D biosystems 248-BD-025), 200 mM cAMP (Sigma D0627), and 200 μM ascorbic acid (Sigma 4034-100). until the generation of d35 CN.

### ESC cell line generation

EHMT1-dTAG-HA-EGFP homozygous targeted, GPI-dTAG-HA-FLAG-EGFP homozygous targeted, and EHMT1-HA-EGFP ZNFxxx-dTAG-V5-mCherry (ZNFxxx-ZNF460, ZNF462, ZNF644) dual homozygous targeted lines were generated in MEL-1 human ESCs using CRISPR-Cas9 engineering as previously described^75^ Homology arms at C terminus of all targeted loci were amplified from the parental gDNA with upstream and downstream base pairs as follows – EHMT1: 602 bp, 550 bp, GPI: 618 bp, 511 bp, ZNF460: 544 bp, 430 bp, ZNF462: 664 bp, 395 bp, ZNF644: 589 bp, 350 bp. We then cloned these PCR products into TOPO vectors and combined them using Gibson cloning to insert FKBP12F36V-2xHA-P2A-NLS-EGFP for EHMT1-dTAG-HA-GFP, 2xHA-P2A-NLS-EGFP for EHMT1-HA-GFP, P2A-FKBP12F36V-HA-FLAG-EGFP for GPI, and FKBP12F36V-2xV5-P2A-mCherry for ZNFxxx targeting. These targeting vectors were transfected together with a pX330-geneticin vector bearing the specific gRNAs targeting the C-terminus of each gene using lipofection in mTeSR media (Stem Cell Technologies, 05850) supplemented with CloneR (Stem Cell Technologies, 05888) solution (1:10). Cells were selected for the presence of pX330 with G418 for 48 h (500 μg/ml – Day 1, 250 μg/ml – Day 2). 2-3 days after selection, cells expressing the EGFP and/or mCherry transcriptional reporter were single-cell sorted by FACS and expanded for characterization. Homozygosity was assessed in individual clones by PCR genotyping and by WB when antibodies were available. The dual-targeted lines were sequentially targeted for EHMT1-HA-GFP, followed by ZNFxxx-dTAG-V5-mCherry.

### Flow cytometry

Single-cell sorting of dTAG clones was conducted with the help of the flow cytometry core facility at Weill Cornell. Live cells were run on the BD Influx, gated for high levels of mCherry and/or EGFP to enrich for homozygosity, and seeded in 96-well plates. After growing, single-cell-derived clones were characterized by live-cell flow. Cells were resuspended in 3 µg/ml DAPI and filtered through a 35 µm cell strainer (VWR, 352235) into FACS tubes. We determined expression of endogenous loci bearing mCherry using the 561 nm (610/20) channel, and EGFP using the 488 nm (530/30) channel. Analysis was performed on a BD Fortessa flow cytometer using FlowJo v10.10 software.

To assess HA, V5, PAX6, or H3K9me2 depletion after dTAG treatment, we assessed protein levels by intracellular flow cytometry. Briefly, 0.75-1×10^6^ cells were collected in a single-cell suspension and processed using an intracellular staining kit (Thermo Fisher, Cat. 00-5523-00). Briefly, cells were fixed for 25 min followed by incubation with 100 µl of 1:800 HA primary antibody (CST-3724S) or 1:500 Alexa Fluor 647 (AF647) conjugate HA antibody (CST-37297) for HA flow; 1:500 AF647 conjugate V5 antibody (CST-D3H8Q), 1:200 AF 647 Mouse anti-Human PAX6 (BD Biosciences,562249), or 1:400 H3K9me2 antibody (ab1220) diluted in permeabilization buffer for 40 min (in dark for conjugated antibody). Primary antibody-incubated cells were then incubated with 100 µl of 1:500 secondary antibody (647DaRb TF A-31573, 555DaRb A-31572, 647DaMs A-31571, 555DaMs A-21202) for 30 min in the dark. Finally, cells were resuspended in 300nM DAPI and filtered through a 35 µm cell strainer (VWR, 352235) into FACS tubes for analysis. Analysis was performed on a BD Fortessa flow cytometer using FlowJo v10.10 software.

### Immunofluorescence

Immunofluorescence samples were fixed on tissue culture plates by aspirating the media and washing with PBS, followed by 10min fixation in 4% paraformaldehyde (PFA) in the dark. Note that d10 NE and d20 NPC samples were collected as a single-cell suspension and replated into Matrigel-coated plates at 1:10 for 2 h to reduce confluence. Cells were then either stored at 4C for up to two weeks or permeabilized using 0.5% Triton X100 for 5 minutes followed by incubation in blocking buffer – 5% goat serum (TF, 16-210-064) and/or donkey serum (Sigma, D9663) depending on the secondary antibody, 0.2% tween-20 and 0.2% fish skin gelatin (Sigma, G7041) diluted in 0.2% Tween-20 in PBS (PBS-T) for 30min. Samples were then incubated with primary antibodies diluted in blocking buffer for 1 h for the respective assays: 1:500 HA (CST-3724S) to validate dTAG-mediated EHMT1 depletion; 1:50 OCT4 (sc5279); 1:50 PAX6 (ab42-6600); 1:200 FOXG1 (E2W8P); 1:2000 MAP2 (ab5392). Primary antibodies were washed out three times with PBS-T at 5min intervals, followed by 5min of blocking buffer and 1 h incubation in the dark with a 1:500 secondary antibody, depending on the primary antibody (647DaRb TF A-31573, 555DaRb A-31572, 647DaMs A-31571, 555DaMs A-21202, 647GaRb A-21245). Cells were washed three times with PBS-T at 5min intervals, then stained with 300nM DAPI for 5min. Images were acquired on an EVOS Cell Imaging System (Thermo Fisher) or an LSM880 confocal microscope (Zeiss). For EHMT1 depletion kinetics, mean signal intensity was secondary control-subtracted and normalized to DMSO for each cell type.

### RNA isolation

Total RNA from cells was extracted using TRIzol (Invitrogen 15596018) and purified using the RNA Clean and Concentrator kit (Zymo Research ZR1014) with DNase treatment. RNA quality and quantity were checked before assays using a NanoDrop or Bioanalyzer.

### RT-qPCR

Extracted RNA was reverse transcribed into cDNA using the iScript kit (Bio-Rad, 1708841). qPCR was performed in triplicate using PowerUp SYBR Green PCR master mix (Thermo Fisher A25778) on an Applied Biosystems QuantStudio 3. Primers spanning exon-intron overlaps are shown in **Supplemental Table 1**.

### RNA-sequencing

Extracted RNA for 48 h dTAG treatment for ES and d10 NE samples, ES◊d3 dTAG samples for EHMT1 and ZNF462, and ES → NPC dTAG for EHMT1 and ZNF462 was processed by the Weill Cornell Genomics Core. Library prep was conducted using the Illumina Stranded Total RNA Prep with Ribo-Zero Plus (Illumina 20040529), and sequencing was performed on an Illumina NovaSeq flow cell with PE 100 bp reads. Extracted RNA for 48 h dTAG treatment for GPI samples was processed by the Weill Cornell Genomics Core. Library prep was conducted using TruSeq Stranded mRNA Library Preparation, and sequencing was performed on a NovaSeq 6000 with PE 100 bp reads. Extracted RNA from NPC and CN samples after 48 h dTAG treatment was processed by Novogene. Library prep was conducted using Directional library preparation (rRNA removal) and sequenced on a NovaSeq X Plus (PE150).

Extracted RNA for single-cell validation at d35 CN was processed by Plasmidsaurus. Samples were prepared by Plasmidsaurus by first generating cDNA using a poly(dT)VN primer. Single-strand sequencing was performed using 3′-end counting with UMIs for deduplication and unique dual indices (UDIs) to prevent index hopping, with a read length of about 90 bp.

### Histone mass spectrometry

Histones were extracted from frozen pellets using a commercial histone extraction kit following the manufacturer’s protocol (Abcam, ab113476). In the subsequent steps, histones were processed as previously described^76^. Briefly, soluble histones were precipitated overnight with 33% cold trichloroacetic acid (TCA, Sigma-Aldrich, T0699-100ml) under rotation at 4°C. Samples were spun at 3400 RCF for 5 min; the supernatant was removed, and histone pellets were rinsed with ice-cold acetone + 0.1% HCl and with 100% ice-cold acetone. Histone pellets were allowed to dry at room temperature. Next, the pellet was dissolved in 50 mM ammonium bicarbonate (pH 8.0), and histones were derivatized using 5 µL propionic anhydride and 14 µL ammonium hydroxide (both Sigma-Aldrich) to adjust the pH to 8.0. The mixture was incubated for 15 min, and the procedure was repeated. Histones were then digested with 1 µg of sequencing-grade trypsin (Promega V5280) diluted in 50 mM ammonium bicarbonate (1:20, enzyme: sample) overnight at room temperature. The derivatization reaction was repeated to derivatize peptide N-termini. The samples were dried in a vacuum centrifuge. Prior to mass spectrometry analysis, samples were desalted using a 96-well plate filter (Orochem) packed with 1 mg of Oasis HLB C-18 resin (Waters). Briefly, the samples were resuspended in 100 µl of 0.1% TFA and loaded onto the HLB resin, which had been equilibrated with 100 µl of the same buffer. After washing with 100 µl of 0.1% TFA, the samples were eluted with a buffer containing 70 µl of 60% acetonitrile and 0.1% TFA and then dried in a vacuum centrifuge.

Samples were resuspended in 10 µl of 0.1% TFA and loaded onto a Dionex RSLC Ultimate 300 (Thermo Scientific), coupled online with an Orbitrap Fusion Lumos (Thermo Scientific). Chromatographic separation was performed with a two-column system, consisting of a C-18 trap cartridge (300 µm ID, 5 mm length) and a picofrit analytical column (75 µm ID, 25 cm length) packed in-house with reversed-phase Repro-Sil Pur C18-AQ 3 µm resin. Peptides were separated using a 30 min gradient from 1-30% buffer B (buffer A: 0.1% formic acid, buffer B: 80% acetonitrile + 0.1% formic acid) at a flow rate of 300 nl/min. The mass spectrometer was set to acquire spectra in a data-independent acquisition (DIA) mode. Briefly, the full MS scan was set to 300-1100 m/z in the Orbitrap with a resolution of 120,000 (at 200 m/z) and an AGC target of 5×10e5. MS/MS was performed on the Orbitrap with sequential isolation windows of 50 m/z, an AGC target of 2×10e5, and an HCD collision energy of 30.

Histone peptide raw files were imported into EpiProfile 2.0 software^77^. From the extracted ion chromatogram, we obtained the area under the curve and used it to estimate the abundance of each peptide. In order to achieve the relative abundance of post-translational modifications (PTMs), the sum of all different modified forms of a histone peptide was considered as 100% and the area of the particular peptide was divided by the total area for that histone peptide in all of its modified forms. The relative ratio of two isobaric forms was estimated by averaging the ratio for each fragment ion with different mass between the two species. The resulting peptide lists from EpiProfile were exported to Microsoft Excel and further processed for detailed analysis.

### Immunoprecipitation mass spectrometry

Cells were collected for immunoprecipitation mass spectrometry (IP-MS) using Accutase, and nuclear lysate was collected using the NE-PER kit (TF 78833). 100 µg nuclear lysate was added to 25 µl magnetic anti-HA beads (TF 88836), which had been washed twice with TBS with 0.05% Tween-20 (TBS-T). Samples were rotated at room temperature for 1 h, then washed twice with TBS-T, followed by ultrapure water. On-bead treatment and trypsin digestion were performed with all dilutions in 50 mM pH 8.0 Ammonium bicarbonate as follows: 25 µl solution of 5mM DTT was added to beads for 45 min at room temperature, followed by 2.5 µL 200 nM iodoacetamide for 30 min in the dark, after which 500 ng of sequencing-grade trypsin (1:20, enzyme: sample) was added for overnight digestion at room temperature. Peptide concentrations were quantified using the Pierce Quantitative Colorimetric Peptide Assay (Thermo Scientific 23275). Prior to mass spectrometry analysis, samples were desalted using a 96-well plate filter (Orochem) packed with 1 mg of Oasis HLB C-18 resin (Waters). Briefly, the samples were resuspended in 100 µl of 0.1% TFA and loaded onto the HLB resin, which had been equilibrated with 100 µl of the same buffer. After washing with 100 µl of 0.1% TFA, the samples were eluted with a buffer containing 70 µl of 60% acetonitrile and 0.1% TFA and then dried in a vacuum centrifuge. Samples were resuspended in 10 µl of 0.1% TFA and loaded onto a Dionex RSLC Ultimate 300 (Thermo Scientific), coupled online with an Orbitrap Fusion Lumos (Thermo Scientific). Chromatographic separation was performed with a two-column system, consisting of a C-18 trap cartridge (300 µm ID, 5 mm length) and a picofrit analytical column (75 µm ID, 25 cm length) packed in-house with reversed-phase Repro-Sil Pur C18-AQ 3 µm resin. Peptides were separated using a 90 min gradient from 4-30% buffer B (buffer A: 0.1% formic acid, buffer B: 80% acetonitrile + 0.1% formic acid) at a flow rate of 300 nl/min. The mass spectrometer was set to acquire spectra in a data-dependent acquisition (DDA) mode. Briefly, the full MS scan was set to 300-1200 m/z in the Orbitrap with a resolution of 120,000 (at 200 m/z) and an AGC target of 5×10e5. MS/MS was performed in the ion trap using the top speed mode (2 secs), an AGC target of 1×10e4, and an HCD collision energy of 35.

Proteome raw files were searched using Proteome Discoverer software (v2.5, Thermo Scientific) with the SEQUEST search engine and the SwissProt human database (updated April 2023). The total proteome search included variable modification of N-terminal acetylation and fixed modification of carbamidomethyl cysteine. Trypsin was specified as the digestive enzyme with up to 2 missed cleavages allowed. Mass tolerance was set to 10 pm for precursor ions and 0.2 Da for product ions. Peptide and protein false discovery rate was set to 1%. Following the search, data were processed as previously described^78^. Briefly, proteins were log2-transformed, normalized to each sample’s mean, and missing values were imputed using a normal distribution centered at the sample mean with a standard deviation of 2. Statistical regulation was assessed using a heteroscedastic T-test (if p-value < 0.05). Data distribution was assumed to be normal, but this was not formally tested.

### ChIP sequencing

ChIP-seq was performed as previously described^79^ with some modifications. For EHMT1 ChIP, 30 million HA-tagged cells were double-crosslinked by first incubating with 2 mM DSG (Sigma 80424-5MG-F) for 50 min, followed by 1% formaldehyde at RT for 10 min, and then quenched with 125 mM glycine for 5 min at RT. For H3K27ac, ZNF462 and ZNF644 ChIP, 15 million cells were single-crosslinked by incubating with 1% formaldehyde at RT for 10 min, and then quenched with 125 mM glycine for 5 min at RT. Cells were then flash frozen and stored at –80 before further processing. Cells were resuspended at 15 million per 300ul lysis buffer (10 mM Tris, pH 8, 1 mM EDTA, 0.5% SDS) and sonicated in a Bioruptor Pico for 15-30 cycles and then centrifuged for 10 min at 4 °C at 17,000 × *g*. 5% of the sample was first checked for appropriate sonication by DNase treatment at 65C for 2 h, followed by running on a 1% agarose gel to confirm 200-700bp fragments. Supernatants were precleared for 1 h with 20 μL of Protein A Dynabeads (Thermo Scientific 10-001-D) per sample. 5% of each sample was removed and frozen as an input sample. Samples were diluted 5 times with dilution buffer (0.01% SDS, 1.1% Triton,1.2 mM EDTA,16.7 mM Tris, pH 8, 167 mM NaCl) and incubated with HA antibody (5 μg/30 million starting cells) (CST 3724), V5 antibody (5ug/30mil starting cells) (CST 13202), or H3K27ac antibody (4.5ug/15mil starting cells) (AM 91193) overnight with rotation at 4 °C. The next day, protein A Dynabeads pre-blocked with 1 mg/mL BSA protein were added to each sample (30 μL Dynabeads per sample). Samples were incubated for 3.5 h at 4 °C. The beads were immobilized on a magnet and washed twice with low-salt buffer (0.1% SDS, 1% Triton X-100, 2 mM EDTA, 150 mM NaCl and 20 mM Tris pH 8), twice in high-salt buffer (0.1% SDS, 1% Triton X-100, 2 mM EDTA, 500 mM NaCl and 20 mM Tris pH 8), twice in LiCl buffer (0.25 M LiCl, 1% NP-40, 1% deoxycholic acid (sodium salt), 1 mM EDTA and 10 mM Tris pH 8) and once in TE buffer. DNA elution from the beads was performed by incubation with 125 µl of elution buffer (1% SDS and 10 mM NaHCO3) for 30 min at 65 °C and 2000 rpm in a thermomixer. The supernatants and inputs were reverse-crosslinked by overnight incubation at 65 °C with 100 µg/ml proteinase K. After treatment with RNase A for 30 min at 37 °C, the DNA was purified using a DNA cleanup and concentration kit (Zymo, zd4014). The immunoprecipitated DNA was used for library preparation using the Thru-plex kit (Takara R400675) with Thru-plex indexes (Takara R400665-8) and sequenced on Illumina NovaSeq flow cells PE100.

### ULI-NChIP

A total of 100,000 cells were used as input for H3K9me2 ChIP, with two independent replicates. ULI-NChIP was conducted following a published procedure^28^. Briefly, flash-frozen cell pellets were resuspended in Sigma EZ Nuclei Isolation Lysis Buffer (NUC101) containing 1× protease inhibitor cocktail (Roche 04693132001) and 1 mM PMSF (TF 36978), and then digested with MNase (NEB M0247) and spiked in with ∼3% mouse ESCs for normalization. The digested chromatin was rotated for 1 h at 4 °C, then pre-cleared with Protein A/G beads (Dynabeads, Life Technologies #1006D), and 10% was saved as an input control. In total, 1 μg per 100k cells of H3K9me2 antibody (ab1220) was added to Protein A/G beads for 3 h at 4 °C to form the antibody complex. Precleared chromatin was added to the antibody complex and rotated at 4 °C overnight. The antibody-bound chromatin was then eluted from the magnetic beads using 100 mM NaHCO3 and 1% SDS at 65 °C for 1 h and purified alongside the input chromatin. We prepared libraries from the eluted and input chromatin using ThruPLEX DNA-Seq (Takara R400675) following the manufacturer’s instructions. Libraries were sequenced on an Illumina NovaSeq X Plus platform with PE100 reads.

### ATAC-seq

ATAC-seq was performed on MEL-1 OTX2-HA-dTAG cells^75^ following a previously published protocol^80^. We washed 50,000 cells per replicate in 1x PBS and resuspended them in 50 µl ATAC-resuspension buffer (10 mM Tris-HCl pH 7.4, 10 mM NaCl, 3 mM MgCl2) supplemented with 0.1% (v/v) IGEPAL CA-630, 0.1% Tween-20, and 0.01% Digitonin for nuclei isolation. Samples were incubated on ice for 3 min, spun down, and washed with 1 mL of cold ATAC-resuspension buffer containing 0.1% Tween-20. Nuclei were pelleted at 500 rcf for 10 min at 4 °C. After careful aspiration of the supernatant, the cell pellet was resuspended in 50 µl of transposition mix (25 µl TD buffer, 0.8 ul homemade Tn5, 18.2 µl 1x PBS, 0.5 µl 1% digitonin, 0.5 µl 10% Tween-20, 5 µl H2O) by pipetting up and down 6 times. The reaction was incubated at 37 °C for 30 min in a thermomixer at 1000 rpm. DNA was isolated using a commercial kit (Zymo, zd4014). Libraries were generated using the NEBNext HiFi 2x PCR master mix (NEB, #M0541), with each sample having a unique barcode and a universal primer, followed by 10 cycles of amplification. Finally, samples underwent a double-size selection (0.5x-1.3x) with SPRIselect beads (Beckman Colter, B23317) and were sent for TapeStation quality check. Libraries were sequenced on an Illumina NovaSeq X Plus 1.5B/10B Flow Cell (PE50).

### CUT&RUN

d3 NE CUT&RUN was performed on MEL-1 OTX2-HA-dTAG cells following previously published protocols^75^. Cells were dissociated into a single-cell suspension using Accutase, and 500,000 cells were used per replicate. BioMag Plus Concanavalin A beads (ConA beads, PolySciences NC1358578) were washed twice in 100μl cold Bead Activation Buffer (20mM HEPES pH 7.5, 10mM KCl, 1mM CaCl2, 1mM MnCl2). 20μL of ConA beads per CUT&RUN reaction were used. Sorted cells were washed twice in Wash Buffer (20mM HEPES pH 7.5, 150mM NaCl, 0.5 mM spermidine (Acros AC132740050), ½ Protease Inhibitor tablet) and bound to activated ConA beads in Wash Buffer for 10 minutes at room temperature. Samples were incubated in PCR strip tubes with 1:100 anti-HA (CST C29F4 #3724) in Antibody Buffer (Wash Buffer with 2mM EDTA and 0.01% Digitonin (Millipore Sigma 300410)) overnight with gentle rocking at 4°C. Samples were washed twice in Digitonin Buffer (Wash Buffer with 0.01% Digitonin), and Protein A/G MNase (EpiCypher 15-1016) was bound to the samples for 1 hour at 4°C. ProteinA/G MNase was diluted to 20x in 50μl Antibody Buffer according to the manufacturer’s instructions. Samples were washed twice in Digitonin Buffer, and MNase digestion was initiated with 1 μl of 100 mM CaCl2 in 50 μl of Digitonin Buffer for 2 hours. To extract bound chromatin fragments, digestion was quenched using 33μl STOP Buffer (340mM NaCl, 20mM EDTA, 4mM EGTA, 50μg/ml RNAse A, 50μg/ml glycogen, 0.015 ng/μl E. coli spike-in), and samples were incubated at 37°C for 20 minutes. Chromatin was separated from ConA beads by centrifuging for 5 minutes at 16,000g and saving the supernatant. DNA was purified by adding 0.1% SDS and 5 μg Proteinase K for 10 min at 70°C, performing phenol-chloroform extraction, and precipitating in 100% ethanol at –80°C overnight. Pellets were then washed in 100% ethanol and resuspended in 12 μl nuclease-free water. CUT&RUN libraries were prepared using the Thruplex Library Prep kit (Takara R400676) according to the manufacturer’s instructions. The number of cycles for library PCR amplification was determined by running 10% of each library sample on an Applied Biosystems qPCR machine with 0.25 μl Eva Green dye (Biotium 31000) for 40 cycles, then adding cycles to each sample. Libraries were size-selected with 1.5x volume SPRI beads (Beckman Coulter B23317) and sequenced at the Weill Cornell Genomics Resources Core Facility on an Illumina NovaSeq 6000 (PE-50, 30 million reads per sample).

### Single cell RNA-seq

Neuronal cells at d35 were washed thrice in PBS with 100 mg/ml DNAseI, and incubated with Accutase. Cells at indicated stages were dissociated using Accutase supplemented with Neuron Isolation Enzyme for Pierce (Thermo 88285) solution at 1:50 at 37 °C for 45–60 min and gently dissociated to single-cell suspensions via pipetting. After washing in PBS, single-cell suspensions were counted and resuspended to 1 million cells in 50 µl of TruStain FcX blocking reagent (BioLegend 422301) diluted 1:10 in FACS buffer and incubated at 4 °C for 10 min. Meanwhile, the hashtag antibody pool from TotalSeq-B (BioLegend B362116, B319427, B341071, B341070) was spun down at 14000 RCF at 4 °C for 10 min. We added 0.5 µL of hashtag antibody to each sample for pooling, then incubated the samples for 30 min at 4 °C. After 30 min, we centrifuged the samples for 5min at 400 × g, washed with FACS buffer, and repeated the wash twice. The final cells were counted and pooled to 1500 cells/ul per sample in 1× PBS with 0.04% BSA and 0.2 U μl−1 Ribolock RNAse inhibitor (Thermo EO0382). The Epigenomics Core at Weill Cornell generated single-cell libraries according to the manufacturer’s instructions (10x Genomics Chromium Single Cell Gene Expression – Cell Multiplexing), including antibody capture and gene expression library processing.

### dTAG treatment and washout

For 48 h depletion experiments, 250 nM dTAG^v1^ diluted in DMSO was added at 0 h, and cells were collected at 48 h. For the longer-term depletion experiments, 250 nM dTAG treatment was added when cells were plated at the ES stage (d-1) in E8 Flex media for neural induction at d0. Following this, 10nM dTAG was added every 2 days for the length of each experiment. Note that the same volume of DMSO was added to control cells in all dTAG experiments.

For dTAG washout at d20, cells that had received DMSO or dTAG treatment from d-1 to d20 were collected as a single-cell suspension. Cells were centrifuged at 300g, washed 3 times in prewarmed NB/B27 medium, and then split and plated for induced neurogenesis as above. We split the d-1-20 DMSO samples into DMSO and dTAG media (DMSO, NPC◊CN, respectively), and the d-1-20 dTAG samples into DMSO and dTAG media (ES•NPC, ES◊CN, respectively).

### Data processing

All genomics datasets were processed using custom snakemake (v6.6.1) pipelines^81^ and aligned to the hg38 genome with the Gencode v38 annotation unless otherwise specified. RStudio version 2026.4.0.526 was used for quantitative analysis and figure generation.

### RNA-seq

FASTQ files were processed with STAR-multi alignment as previously established^31,82^. Adapters were trimmed using TrimGalore (v0.6.10) followed by mapping with STAR (v2.7.10)^83^. Counts were obtained using featureCounts (subread v2.1.1) and imported into R. DESeq2 (v1.50.2) was used for paired comparisons and differential gene analysis, with count normalization. See **Supplementary Table 1** for DESeq2 outputs.

We conducted bioinformatic analysis of bulk RNA-seq data for d35 CN single-cell RNA-seq validation using Plasmidsaurus’s workflow. FastQ generation and demuxing were done with BCL Convert v4.3.6 and fqtk v0.3.1. Read filtering was done using FastP v0.24.0: poly-X tail trimming, 3’ quality-based tail trimming, a minimum Phred quality score of 15, and a minimum length requirement of 50 bp. Then, alignment to the hg38 genome was performed using the STAR aligner v2.7 with non-canonical splice junction removal, and unmapped reads were output and BAM files were coordinate-sorted using amtools v1.21. PCR and optical duplicates were removed using UMICollapse v1.1.0, and QC was performed with RustQC v0.2.1. and MultiQC v1.33. Gene– expression quantification was done using featureCounts (subread package v2.1.1) with strand-specific counting, multi-mapping read fractional assignment, exons and three-prime UTR as the feature identifiers, and grouped by gene_id. Final gene counts were annotated with gene biotype and other metadata extracted from the reference GTF file. CPM values were obtained with edgePython v0.2.5 and provided in Excel files. We imported these into R and used them to plot different conditions.

The SFARI gene database, release 04-03-2023, was used to compute the autism gene score. Gene ontology analysis was done using enrichGO^84^ with the Biological Processes ontology against the ‘org.Hs.eg.db’ organism database, using all genes as background for the 48 h dTAG experiment, and against genes with non-NA padj values for either EHMT1 or ZNF462 for the d20 NPC RNA-seq experiment. We then performed K-means clustering and generated heatmaps using the ComplexHeatmap^85^ package. We filtered terms for q-value < 0.05 and presented the selected terms. We defined monotonic up genes as those with cell-type mean DESeq2 counts ES<NE<NPC<CN and count CN > 50, and monotonic down genes as those with cell-type mean DESeq2 counts ES>NE>NPC>CN and count ES> 50. Overlap. Significance of overlap between ES◊NPC differential genes (padj <0.05, |log2FC|>0) and SFARI genes was calculated using a hypergeometric test function where N = all genes, K = Category 1or 2 SFARI genes within all genes, n = genes per DEG category, and k = Category 1 or 2 SFARI genes per DEG category.

### ChIP-seq

ChIP sequencing was analyzed using ‘repeat-aware’ alignment with STAR-multi, as previously described^31^. Alignment was performed using STAR (v.2.7.0), duplicates were marked using Picard (v2.26.10), and removed using Samtools (v1.1.4). Blacklist removal was done using ENCODE hg38 blacklist v2 with Samtools. Peaks were called with p-value < 0.05 using MACS2 (v2.2.6) against matched input samples. BigWig files were generated using deepTools (v3.5.1) bamCoverage with a 10 bp bin size, RPGC normalization, ignoring duplicates and blacklist regions.

BigWig signal for histone datasets at ES DEG categories were imported using rtracklayer^86^ over the span of +/− 1kb from the TSS. The values were then normalized as indicated and plotted with ggplot::violinplot. ChIP bigwig replicate samples were merged using deepTools (v3.5.1) bigWigAverage with average signal for heatmap generation. EHMT1 signal with ZNF462 and ZNF644, DMSO, and dTAG treatment was calculated for EHMT1 peaks using subread featureCount^87^. K-means and heatmap matrix were generated using the SeqPlots R package^88^and plotted using ComplexHeatmap (see **Supplementary Table 3**). EHMT1 peaks for ES and d3 were combined using GRanges^89^. We used ChIPseeker annotations to assign the nearest gene to peaks, defining the promoter region as +/− 2.5kb from the TSS and using the TxDb file from the Gencode v38 annotation. ChromHMM^91^ 18-state model for H1 ESCs was used for annotation. Nearest peak to gene annotation was done manually by defining +/− 2.5kb TSS as TSS, peaks within the gene body as gene body, and using GRanges::distanceToNearest to get the distance between TSS and peak. Genome tracks were plotted using PlotGardener^92^ package in R with custom genome assembly generated for Gencode v38 annotation. To identify transcription factors and chromatin-associated proteins enriched at the tested genomic regions, we performed Locus Overlap Analysis (LOLA) in R^93^ using a custom region database generated from ChIP-Atlas. More than 2,100 hg38-aligned ChIP-seq peak sets corresponding to transcription factors and other chromatin-associated factors profiled in human embryonic stem cells and other pluripotent cell types were downloaded from ChIP-Atlas. Peak files and associated metadata were formatted according to the LOLA region database structure and incorporated as a custom library. Enrichment was assessed using Fisher’s exact test, and P values were corrected for multiple testing using the Benjamini–Hochberg method. K-means clusters were compared against all peaks, and LOLA-enriched ChIP factors were filtered to have adjusted p-values < 0.05 and DESeq2 counts > 50 in either ES or d3 NE.

The following published datasets were used – bigwigs were used as provided for H3K27me3 (GSM8384176)^94^, H3K4me3 (GSM2029350)^95^, LaminB1 (GSM5669226)^96^. Bigwigs files for H3K9me3(GSE159718)^97^ and POU5F1(OCT4, GSM3381968)^43^ were converted using bigwig CrossMap from hg19 to hg38. ESC ATAC (GSE109524)^43^ was processed from FASTQ files for ESC.

### ULI-NChIP-seq

H3K9me2 ChIP sequencing analysis was done as above with a few additions. We obtained mouse spike-in values by aligning to the mm10 genome and divided total BAM coverage by the hg38 genome BAM reads. After generating RPGC-normalized bigwigs, they were re-binned at 10 kb resolution with a 5 kb sliding step using a custom R script, as previously described^31,98^. To identify H3K9me2 domains, regions of the genome with high H3K9me2 signal (defined as log2(ChIP/Input) >75th percentile) were extended up– and downstream until a region with low H3K9me2 enrichment was encountered (defined as log2(ChIP/Input) <20th percentile). The following packages were used: GenomicRanges, plyranges^99^, rtracklayer, and org.Hs.eg.db.

### CUT&RUN

CUT&RUN fastq files were processed as previously described^100^, using a custom snakemake (v6.6.1) pipeline^101^. Briefly, reads were trimmed using bbmap (v38.95) with parameters ktrim=r ref=adapters rcomp=t tpe=t tbo=t hdist=1 mink=11. Trimmed reads were aligned to the hg38 reference genome using Bowtie2^102^ (v2.3.5.1) with parameters –-local –-very-sensitive-local –-no-mixed –-no-discordant –-phred33 –I 5 –X 999. Reads with a MAPQ score less than 5 were removed with samtools^103^ (v1.14). PCR duplicates were removed with Picard (v2.26.10) using parameter VALIDATION_STRINGENCY=LENIENT. Fragments between 20 and 120 bp were isolated using a custom awk script and used for downstream analyses as previously recommended^104^. BigWigs were generated from BAM files using bedtools^105^(v2.30), normalizing to sequencing depth using the formula: (effective_genome_size / read_count ^∗^ read_length) with 1870000000 as the effective genome size. Bam files were converted to bed files with bedtools (v2.30) with the parameter –bedpe. MACS^106^ (v2.2.6) was used to call peaks on individual and merged replicates using parameters –g 1870000000 –-nomodel –-seed 123 –keep-dup all.

### ATAC-seq

Adapters were trimmed off from paired-end reads using bbduk.sh (v2.4.5) with the following option: ktrim=r ref=adapters rcomp=t tpe=t tbo=t hdist=1 mink=11. Filtered reads were aligned using bowtie2^102^ (v2.4.5) with the options –-local –-very-sensitive-local –-no-unal –-no-mixed –-no-discordant –-phred33 –I 10 –X 2000 –t –p 20. Mapped primary alignments were retained using samtools with the filter of –bq 5. PCR duplicates were removed using the MarkDuplicates module from picard (v2.27.4) with the following options: VALIDATION_STRINGENCY=LENIENT REMOVE_DUPLICATES=true ASSUME_SORTED=true. Mitochondrial reads and reads overlapping hg38 blacklist regions were removed using samtools and BEDTools^105^ (v2.30.0). Peaks were called with MACS2 callpeak, with options: –-keep-dup all –-call-summits –nomodel –-shift –100 –-extsize 200 for summits and –-keep-dup all –-broad –-nomodel –-shift –100 –-extsize 200 for broadpeaks. DeepTools bamcoverage was used to generate bigwig files from bams with options: –p 10 –-ignoreForNormalization chrX chrM –-normalizeUsing RPGC –-extendReads 200 –ignoreDuplicates.

### Single cell RNA-seq

Gene expression and antibody-capture feature-barcode matrices was generated by the Epigenomics core at Weill Cornell using Cell Ranger 9.0. Downstream analysis was done using Seurat (v5.5)^107^ in RStudio. Cells were quality-filtered to nCount_RNA > 1000, nFeature_RNA > 1000, percent.mt <20 and demultiplexed with HTODemux after CLR normalization of the HTO assay(positive.quantile = 0.9) and only singlets were retained. Retained cells were then processed using NormalizeData, FindVariableFeatures, ScaleData, RunPCA, FindNeighbors, FindClusters and RunUMAP using default settings. All downstream analysis was visualized on the shared UMAPs generated with all treatment conditions combined. Differentially expressed genes were determined using Seurat FindMarkers (Wilcoxon-rank-sum test, min.pct = 0), comparing treatment conditions separately for each cell type. Significant genes were called with |avg_log2FC| > 0.5 and adjusted p value <0.05. Functional enrichment was done using gProfiler::gost^108^ for Homo sapiens with ordered_query = TRUE (ranked by fold change), and other default settings. UMAPs and violin plots were made using Seurat and SCP package. Relevant enrichment terms were then selected. To compare cell distributions across the UMAP, each condition was sub-sampled to the condition with the lowest cell number and using FNN library to get k nearest neighbor densities (k= 400).

Cell subpopulations were defined using the Leiden algorithm in Scanpy v1.10.3 at resolutions 0.3. Cluster identities were assigned by comparing canonical marker gene expression with the annotated reference datasets of each postmortem reference. This defined cortical projection neurons (CPN, CPN2), D1-, D2– and D1/D2-medium spiny neurons, layer 1 interneurons and glioblasts.

Neuronal maturation and neuronal function were scored per cell with scanpy.tl.score_genes. Two gene signatures were scored per cell with scanpy.tl.score_genes on log-normalized counts: a 106-gene neuronal maturation signature and a 67-gene neuronal function signature^27^. Scores were compared between conditions separately within each cell type using two-sided Mann– Whitney U tests. Each treatment was compared with DMSO; a Bonferroni correction was applied within each comparison family and cell type. Effect sizes are reported as rank-biserial correlation, with positive values indicating a higher score in the second group of the comparison.

Gene regulatory networks were inferred with pySCENIC^61,109^. Prior to analysis, genes were filtered to retain only those present in both the in vitro dataset and the motif ranking feather files used for motif analysis. Co-expression modules were built with GRNBoost2 using the human transcription factor list allTFs_hg38. Modules were pruned by cisTarget motif enrichment against the hg38 motif collection v10; NES ≥ 2.0, minimum 20 genes per regulon, dropout masking enabled, all modules retained, giving 89 activating and 163 repressing regulons across 25,070 cells. Per-cell regulon activity was scored with AUCell. Regulons whose driving transcription factor was detected in fewer than 10% of cells were excluded.

Regulon activity was compared between conditions separately within each of the four most abundant cell types (CPN, D1-MSNs, D2-MSNs, L1 INT), using two-sided Mann–Whitney U tests with Bonferroni correction and rank-biserial effect sizes. Cell-type-specific responses comprise all regulons significantly changed (padj <0.001, effect_r >0.1) between DMSO and dTAG in at least one cell type and were filtered to include regulons containing a DNA-binding domain.

