## Supplemental Figures and legends for "Developmental rewiring of EHMT1 function during human cortical neurogenesis"

### Supplementary figures

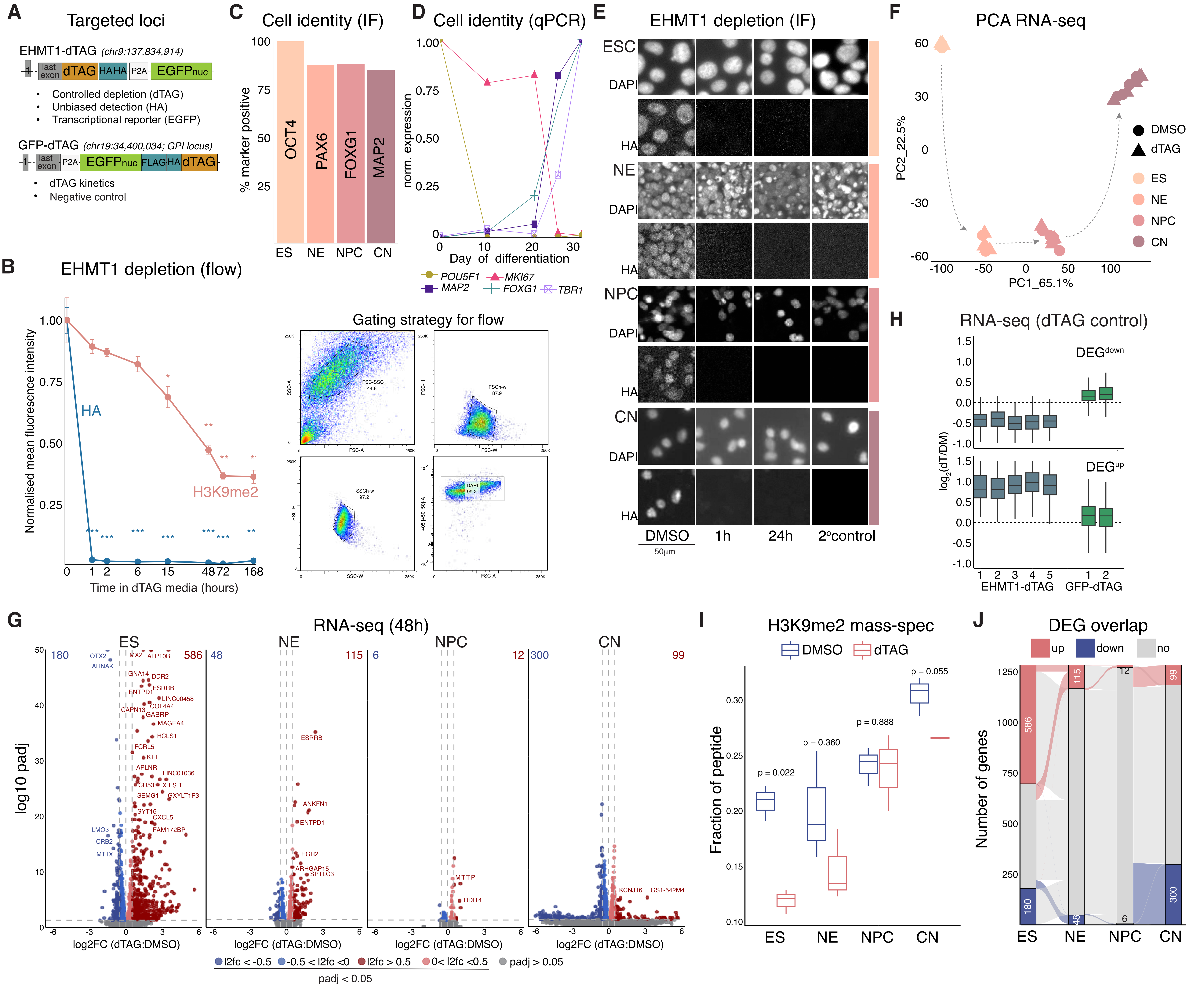


**Figure S1. EHMT1 has distinct stage-specific transcriptional targets during human cortical neurogenesis, related to Figure 1**

A. Targeting strategy and applications for EHMT1-dTAG and GFP-dTAG alleles.
B. Flow cytometry quantification of EHMT1 (HA) and H3K9me2 depletion kinetics. Values were background-subtracted using the secondary-only control and normalized to DMSO levels, unpaired t-test.Gating strategy to obtain single cells free of debris on the right.
C. IF quantification of the percentage of cells positive for stage-specific markers; corresponding to Figure 1B. Data for the differentiation of one representative clone are shown.
D. qPCR of stage-specific markers across neuronal differentiation, normalized to the maximum GAPDH-normalized value.
E. IF images showing EHMT1 depletion at 1 and 24 h of dTAG treatment across the four neurogenesis cell types.
F. PCA of top 5% variable genes for RNA-seq at the different stages of neurogenesis, with the directionality of differentiation indicated.
G. RNA-seq volcano plots following 48 h of dTAG treatment at each stage.
H. RNA-seq fold change of EHMT1-dTAG DEGs after 48 h of dTAG treatment of EHMT1-dTAG and GFP-dTAG ES.
I. Histone mass spectrometry quantification of H3K9me2, expressed as a fraction of total H3K9–H3K17 peptide, following 48 h of dTAG treatment at each stage (paired t-test).
J. Alluvial plot of all DEGs (padj < 0.05, |log2FoldChange| > 0.5) across ES, NE, NPC, CN stages. Number of up- and down-regulated DEGs per cell type indicated.

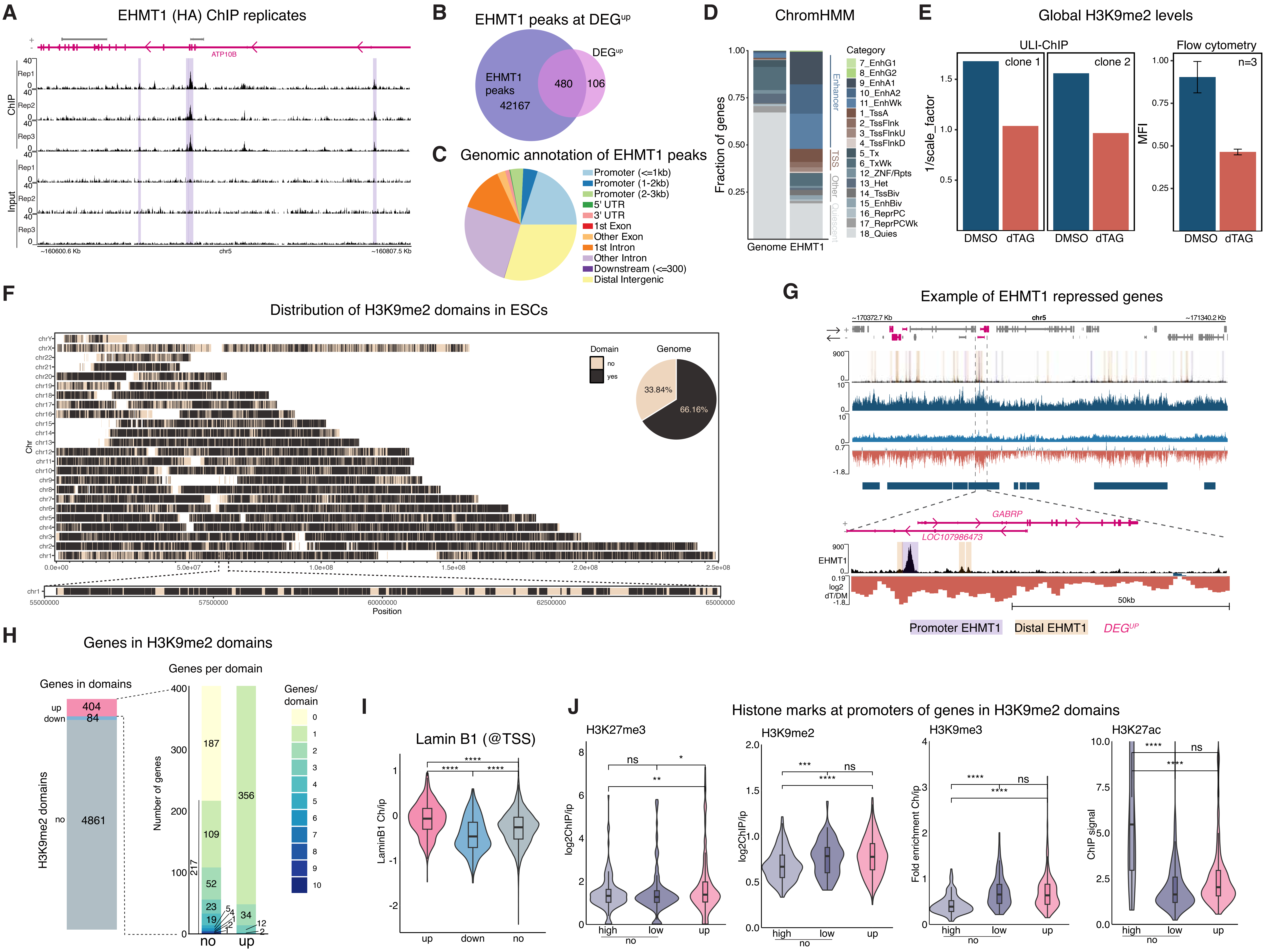


**Figure S2. Molecular determinants of transcriptional sensitivity to EHMT1 loss, related to Figure 2**

A. Representative ChIP-seq genome browser tracks in ES cells showing EHMT1 ChIP and input signal across three biological replicates.

B. Venn diagram showing overlap between ES EHMT1 peaks (ChIPseeker-annotated) and upregulated DEGs from ES EHMT1-dTAG treatment.

C. ChIPseeker annotation of EHMT1 peaks in ES cells.

D. ChromHMM 18-state chromatin model of the ES genome, overlaid with EHMT1 ChIP peaks and four major categories utilized throughout.

E. Global quantification of H3K9me2 ChIP signal, with corresponding flow cytometry quantification of H3K9me2, under EHMT1 DMSO and dTAG treatment.

F. H3K9me2 domains across the linear genome, with a zoom-in on chr1:55,000,000–65,000,000 to offer a representative view at higher resolution. Pie chart showing the percentage of the genome covered by H3K9me2 domains.

G. Representative ChIP-seq genome browser tracks, as in Figure 2A, with a zoom-in for DEGs GABRP and LOC107986473.

H. H3K9me2 domains classified by whether they contain an upregulated DEG, downregulated DEG, or no DEGs ("Genes in domains"; left). For domains containing at least one upregulated DEG, the distribution of domains by the number of upregulated DEGs and non-DEGs is shown.

I. LaminB1 ChIP (GSM5669226) signal over indicated DEG categories, unpaired t-test.

J. Signal strength of the repressive H3K27me3 (GSM8384176) H3K9me2, and H3K9me3 (GSE159718) marks, as well as the active H3K27ac signal at the TSS of the indicated DEG categories, unpaired t-test.

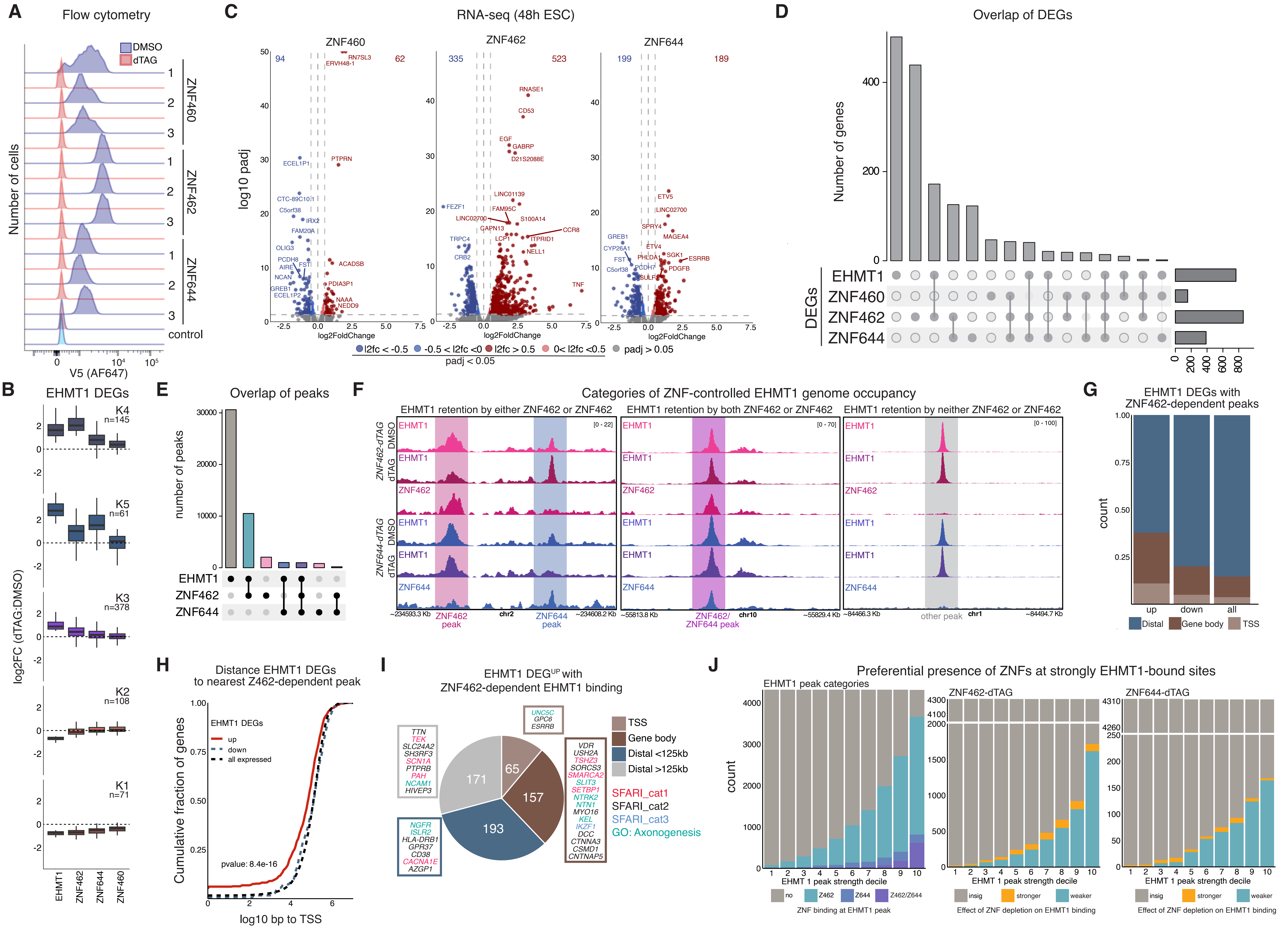


**Figure S3. Target-specific recruitment of EHMT1 by distinct zinc finger proteins, related to Figure 3**

A. Flow cytometry ridge plots of the indicated V5-tagged proteins following 48 h of ES culture in the presence of dTAG or DMSO. Gating strategy as in Fig. S1B.

B. Box plot quantification of the log2FoldChange values of k-means clustered RNA levels shown in Figure 3E.

C. Volcano plots of RNA-seq data following 48 h of dTAG-induced depletion of the respective ZNF proteins.

D. Upset plot showing DEGs shared between depleted proteins (padj < 0.05, llog2FoldChangel > 0.5) following 48 h dTAG treatment.

E. Upset plot showing overlap between EHMT1, ZNF462, and ZNF644 peaks, color-matched to the k-means peak cluster assignments in Fig.3F.

F. Genome tracks of EHMT1 peaks representative for different k-means categories.

G. Bar plot quantifying ZNF462-dependent EHMT1 binding (“Z462” cluster - dashed box in Fig.3F) at loci encoding indicated EHMT1 DEG categories, prioritized by TSS (±2.5 kb), gene body, and distal (>2.5 kb from TSS).

H. Cumulative distribution function showing the distance from the nearest ZNF462-dependent EHMT1 binding site to the TSS of each EHMT1 DEG category. "All expressed" genes were defined as those with a non-NA adjusted p-value (padj) in the EHMT1-dTAG DESeq2 analysis. Kolmogorov–Smirnov test p-value is shown.

I. Genomic distribution of ZNF462-dependent EHMT1 peaks near genes upregulated upon EHMT1 loss using a prioritized hierarchy: TSS (±2.5 kb of the TSS), gene body, <125 kb from the TSS, and >125 kb from the TSS. SFARI genes and genes annotated to the axonogenesis GO term (GO:0007409) are highlighted.

J. EHMT1 peaks split into deciles by peak strength and colored by k-means cluster assignment (left); DESeq2 peak strength for ZNF462-dTAG/ZNF462-DMSO (padj < 0.05, llog2FoldChangel > 0.5) (middle); and DESeq2 peak strength for ZNF644-dTAG/ZNF644-DMSO (padj < 0.05, llog2FoldChangel > 0.5) (right).

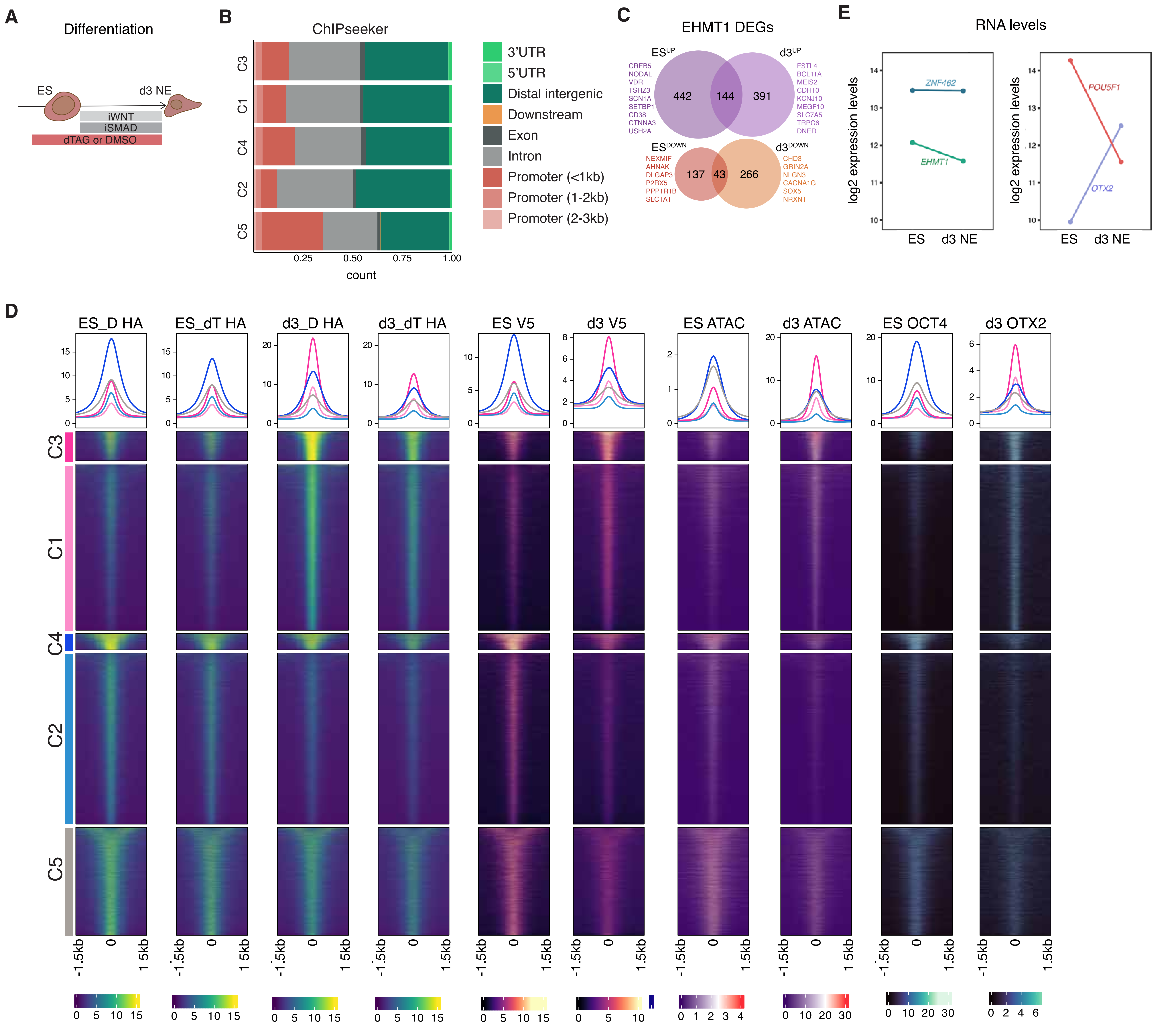


**Figure S4. ZNF462 drives developmental rewiring of EHMT1 genome occupancy during neural induction, related to Figure 4**

A. Schematic representation of the strategy to study the consequences of EHMT1 or ZNF462 depletion during early neural induction.
B. ChIPseeker annotation of ChIP k-means clusters (see Fig.4A).
C. Venn diagram of DEG (padj < 0.05, |log2FoldChange| > 0.5) overlap following EHMT1 dTAG depletion in ES (48 h) and d3 NE (d-1 ES to d3 NE).
D. Combined heatmap of binding profiles for indicated chromatin features at each k-means cluster (see Fig.4A).
E. Expression levels (normalized RNA-seq counts) of the indicate genes in ES and d3 NE.

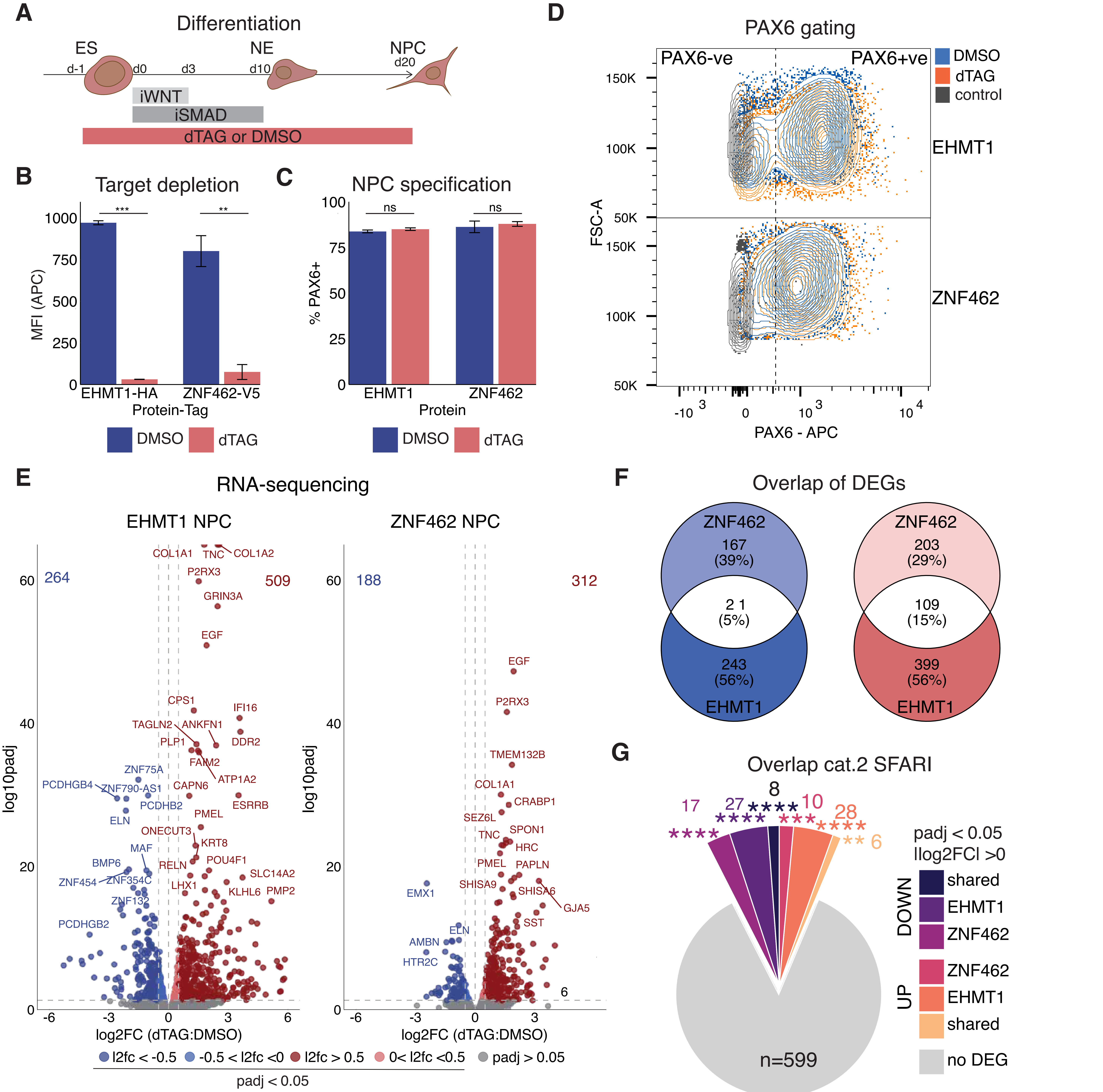


**Figure S5. ZNF462-dependent recruitment underlies shared and distinct EHMT1 transcriptional programs during NPC specification, related to Figure 5**

A. Schematic representation of the strategy to compare the consequences of EHMT1 or ZNF462 depletion throughout NPC specification.
B. Flow cytometry quantification of the levels of tagged proteins at the NPC stage following culture in DMSO or dTAG medium from ES onwards (n = 3 clones), unpaired t-test. Gating strategy as in Fig. S1B.
C. Quantification of the percentage of cells positive for the NPC marker PAX6 following constitutive depletion of the indicated protein.
D. Flow cytometry contour plots showing the PAX6-positive fraction under constitutive depletion of the indicated proteins. Control = cells stained with only secondary antibody.
E. Volcano plots showing RNA-seq gene expression changes triggered by depletion of the indicated protein from ES to d20 NPC.
F. Venn diagram showing the overlap of DEGs identified upon EHMT1 and ZNF462 depletion.
G. Pie chart showing overlap between Category 2 SFARI genes and genes affected by prolonged (ES to NPC) EHMT1 or ZNF462 depletion (padj < 0.05, |log2FoldChange| > 0). P-values calculated as in Fig.5D.

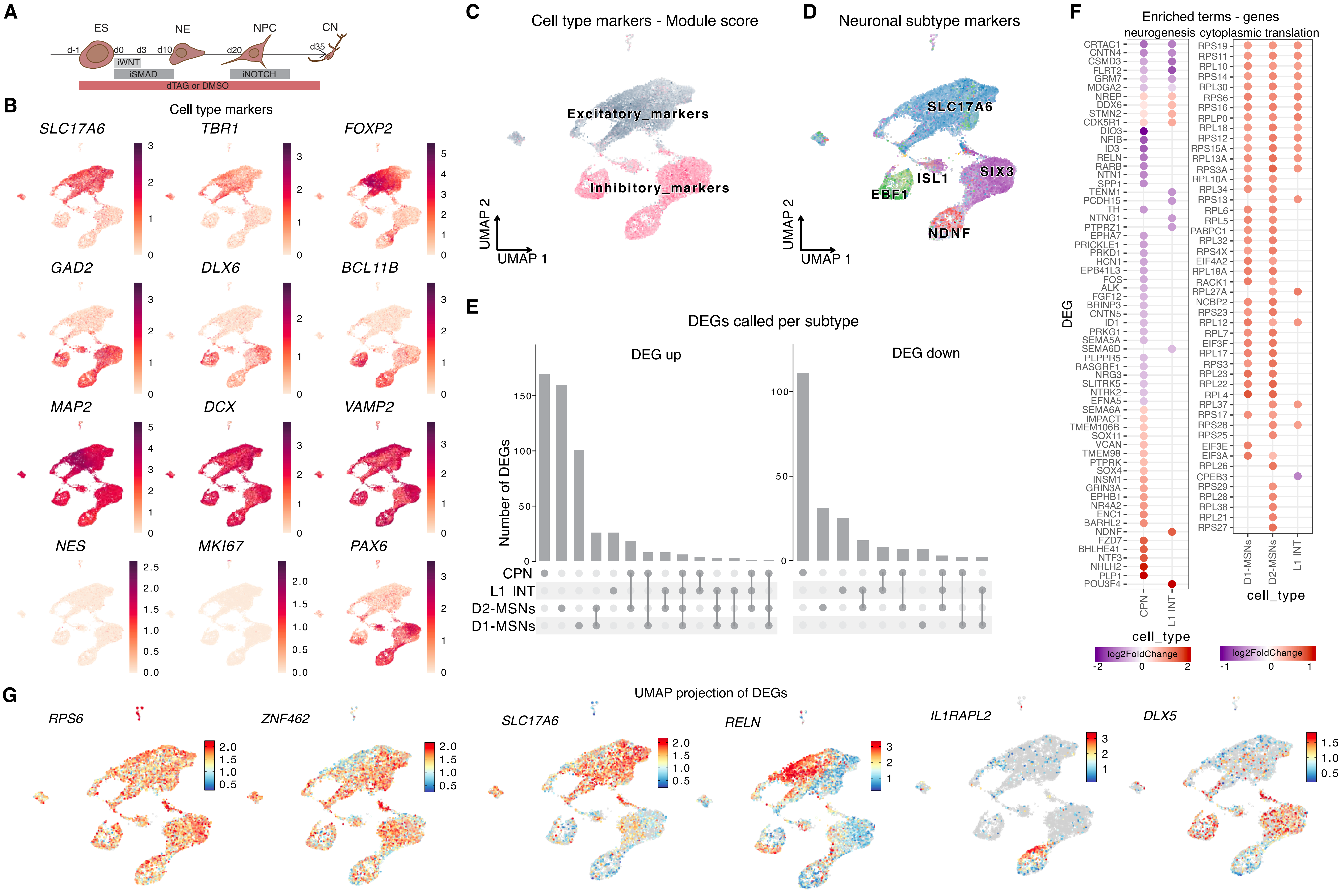


**Figure S6. Developmental EHMT1 deficiency affects cortical neurons in a subtype-specific manner, related to Figure 6**

A. Schematic of the experimental strategy to study the consequences of EHMT1 depletion from the ES to d35 CN stage.
B. UMAP feature plots of cell-type marker genes for glutamatergic (excitatory) cells, GABAergic (inhibitory) cells, pan-neuronal cells and neuronal progenitors.
C. UMAP of Seurat module scores for cortical excitatory (glutamatergic) and inhibitory (GABAergic) cell-type markers. Excitatory markers: SLC17A6, TBR1, FOXP2, CACNA2D1, NRG3 ; inhibitory markers: GAD2, DLX6, SLC32A1, CACNA2D3, BCL11B.
D. Combined UMAP showing expression of a subset of the markers used to assign neuronal identities in Figure 6A.
E. Overlap of genes affected by ES→CN EHMT1 depletion (padj <0.05, Ilog2FoldChangeI > 0.5, up/down) in the different indicated neuronal subtypes.
F. DEGs belonging to g:Profiler functional enrichment terms shared between neuronal subtypes. scRNA-seq expression change (log2FC dTAG/DMSO) indicated.
G. UMAP distributions of Seurat-normalized expression for selected ES→CN DEGs.

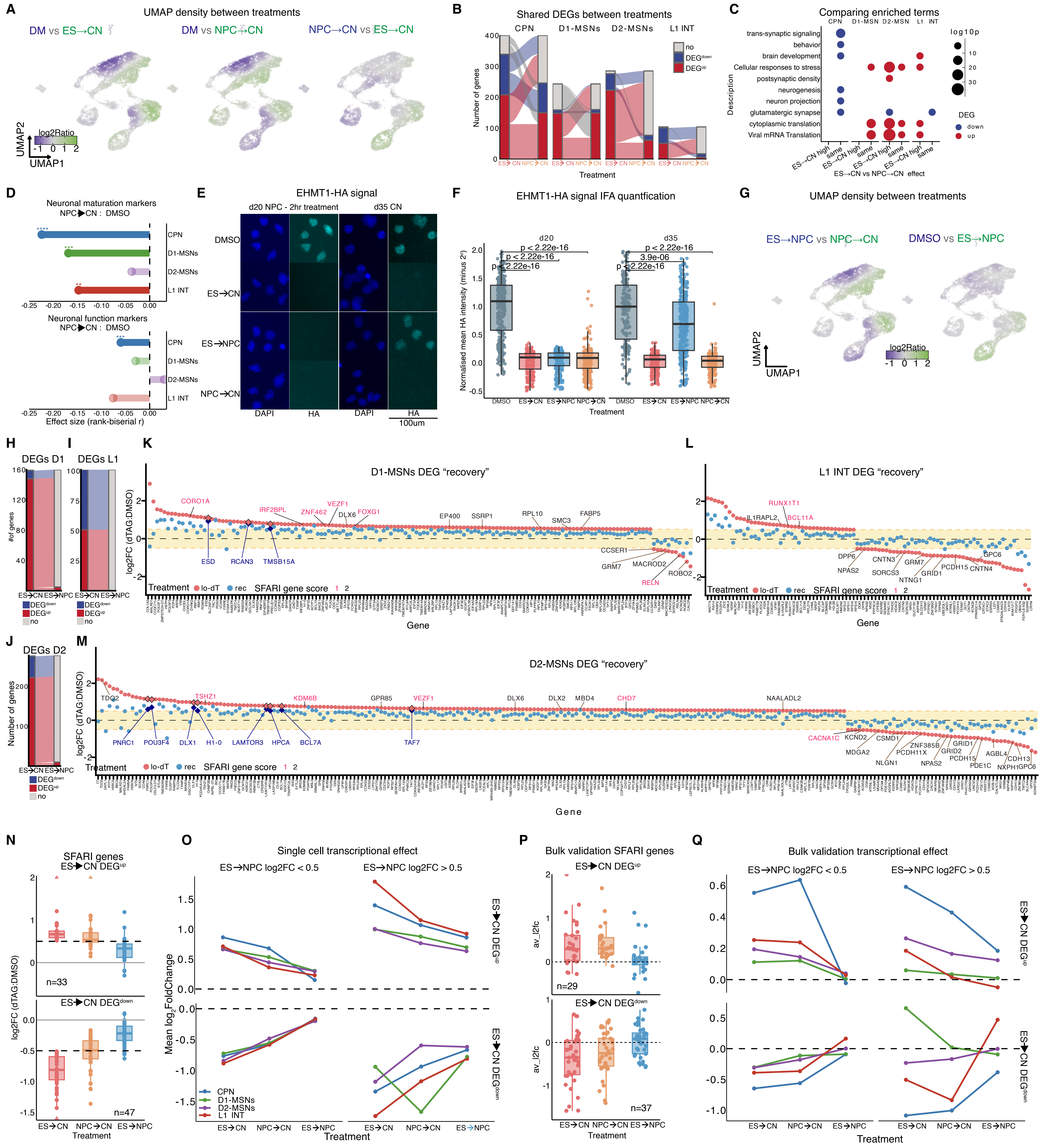

**Figure S7. Accumulation and reversibility of transcriptional dysregulation caused by developmental deficiency of EHMT1, related to Figure 7**

A. D35 scRNA-seq UMAP relative density plots between the indicated treatments. See methods for details.
B. Alluvial plot of ES→CN and NPC→CN DEGs (padj <0.05, Ilog2FoldChangeI > 0.5) in the indicated neuronal subtypes. Color indicates DEG status.
C. Functional terms (gProfiler) enriched in the gene categories identified in Fig. 7C: abs(log2FoldChange(ES →CN - NPC→CN)) >0.3 (ES→CN high), <-0.3(NPC→ CN high),) and -0.3-0.3 (same).
D. Effect of NPC→CN depletion on maturation and neuronal function score as in Fig. 6H,I.
E. Representative IF images of EHMT1-HA signal at NPC and CN stages under indicated treatment conditions.
F. Box plot quantifying EHMT1-HA signal (about 100 cells were assayed per condition). Wilcoxon test p-values are indicated.
G. D35 scRNA-seq UMAP relative density plots between the indicated treatments. See methods for details.
H. Alluvial plot showing ES→CN and ES→NPC DEGs (padj <0.05, Ilog2FoldChangeI > 0.5) in D1-MSN. Color indicates DEG status.
I. As in H, for L1 INT subtype.
J. As in H, for D2-MSNs subtype.
K. Dot plot comparing RNA expression level changes of ES→CN DEGs in D1-MSNs after EHMT1 depletion from ES→CN to ES→NPC. Genes that remain significantly affected upon partial EHMT1 recovery are highlighted with dark blue diamond. Category1 and 2 SFARI genes are highlighted in red and black respectively.
L. As in K, for L1 INT subtype.
M. As in K, for D2-MSNs subtype.
N. RNA expression changes upon indicated timing of EHMT1 depletion for ES→CN DEGs that are Category 1 or 2 SFARI genes .
O. Line plots of the mean log2FoldChange for each ES →CN DEG category per subtype. "ES → NPC log2fc" denotes ES → NPC dTAG Ilog2FoldChangeI irrespective of p-value.
P. Box plots of log2FoldChange in bulk RNA-seq with the same treatment conditions as for the scRNA-seq. Genes and categories as in N.
Q. Line plots of the mean log2FoldChange by bulk RNAseq for genes and categories in O.
